# Equine synovial fluid small non-coding RNA signatures in early osteoarthritis

**DOI:** 10.1101/2020.05.01.066027

**Authors:** Catarina Castanheira, Panagiotis Balaskas, Charlotte Falls, Yalda Ashraf-Kharaz, Peter Clegg, Kim Burke, Yongxiang Fang, Philip Dyer, Tim JM Welting, Mandy J Peffers

**Affiliations:** Department of Musculoskeletal and Ageing Science, Institute of Life Course and Medical Sciences, William Henry Duncan Building, 6 West Derby Street, Liverpool, L7 8TX, UK; Institute of Veterinary Science, University of Liverpool, Chester High Road, Neston, CH64 7TE; Centre for Genomic Research, Institute of Integrative Biology, Biosciences Building, Crown Street, University of Liverpool, Liverpool, L69 7ZB, UK; Institute of Infection and Global Health, University of Liverpool, 8 West Derby Street, Liverpool, L7 3EA, UK; Department of Orthopaedic Surgery, Maastricht University Medical Centre, 6202 AZ Maastricht, The Netherlands

**Keywords:** equine, synovial fluid, osteoarthritis, small non-coding RNAs

## Abstract

**Background:** Osteoarthritis remains one of the greatest causes of morbidity and mortality in the equine population. The inability to detect pre-clinical changes in osteoarthritis has been a significant impediment to the development of effective therapies against this disease. Synovial fluid represents a potential source of disease-specific small non-coding RNAs (sncRNAs) that could aid in the understanding of the pathogenesis of osteoarthritis. We hypothesised that early stages of osteoarthritis would alter the expression of sncRNAs, facilitating the understanding of the underlying pathogenesis and potentially provide early biomarkers.

**Methods:** Small RNA sequencing was performed using synovial fluid from the metacarpophalangeal joints of both control and early osteoarthritic non-Thoroughbred horses. A group of differentially expressed sncRNAs was selected for further validation through qRT-PCR using an independent cohort of synovial fluid samples from control and early osteoarthritic horses. Bioinformatic analysis was performed in order to identify putative targets of the differentially expressed microRNAs and to explore potential associations with specific biological processes.

**Results:** Results revealed 22 differentially expressed sncRNAs including 13 microRNAs; miR-10a, miR-223, let7a, miR-99a, miR-23b, miR-378, miR-143 (and six novel microRNAs), four small nuclear RNAs; U2, U5, U11, U12, three small nucleolar RNAs; U13, snoR38, snord96, and one small cajal body-specific RNA; scarna3. Five sncRNAs were validated; miR-223 was significantly reduced in early OA and miR-23b, let-7a-2, snord96A and snord13 were significantly upregulated. Significant cellular functions deduced by the differentially expressed microRNAs included apoptosis (P < 0.0003), necrosis (P < 0.0009), autophagy (P < 0.0007) and inflammation (P < 0.00001). A conservatively filtered list of 57 messenger RNA targets was obtained; the top biological processes associated were regulation of cell population proliferation (P < 0.000001), cellular response to chemical stimulus (P < 0.000001) and cell surface receptor signalling pathway (P < 0.000001).

**Conclusions:** Synovial fluid sncRNAs can be used as molecular biomarkers for early disease in equine osteoarthritic joints. The biological processes they regulate may play an important role in understanding early osteoarthritis pathogenesis. Characterising these dynamic molecular changes could provide novel insights on the process and mechanism of early osteoarthritis development and is critical for the development of new therapeutic approaches.

## BACKGROUND

Osteoarthritis (OA) remains one of the greatest causes of morbidity and mortality for horses in the UK [1, 2]. Additionally, it is the most common disease affecting the joints in humans, and a significant cause of pain and disability worldwide [3]. This degenerative, age-related joint disease is characterised by a progressive degradation of articular cartilage and concomitant structural and functional change of all joint constituents, including the synovium, the subchondral bone and periarticular tissues [4]. Of multifactorial origin, OA is a product of genetic, mechanical and environmental factors such as age, trauma and occupation [4, 5]. Despite its high prevalence and significant welfare and economic impact, its pathophysiology remains poorly understood and currently available diagnostic tools can only identify the disease when cartilage has already exceeded its capacity for intrinsic repair, and changes can no longer be reversed [6, 7]. As a result, the development of effective treatments is also compromised, and currently recommended therapies are mainly symptomatic.

In the search for molecular biomarkers that could reveal pre-clinical phases of the disease, scientists have focused much of their attention on microRNAs (miRNAs), the best characterised family of small non-coding RNAs. Evolutionarily conserved, these 17-22 nucleotide long molecules regulate gene expression at post-transcriptional level generally by repressing translation or increasing degradation of messenger RNAs (mRNAs). They are involved in different cellular pathways and intercellular communication thus influencing tissue homeostasis [8]. As such, miRNA profiles can be altered as a result of cellular damage and/or tissue injury and altered expression of certain miRNAs is implicated in several diseases, including OA [9–11]. Specific miRNAs have been found to modulate osteoblastogenesis and osteoclastogenesis, chondrogenesis and cartilage degradation, synovial inflammation and neurogenesis, thus contributing to the development and progression of OA; comprehensive reviews of which miRNAs are involved in each of these processes can be found elsewhere [12–14]. miRNAs can be found intracellularly or extracellularly, circulating in virtually any biological fluid in a remarkably stable manner [15–17]. Because biological fluids are generally obtainable through minimally invasive techniques, circulating miRNAs are attractive candidates for disease diagnosis, monitoring and prognostication [18, 19]. Interest in other classes of small non-coding RNAs such as small nucleolar RNAs (snoRNAs) has recently emerged. Mostly known for their housekeeping functions, snoRNAs have canonical roles in the chemical modification of RNA substrates such as ribosomal RNAs, but can also exhibit miRNA-like activity [20]. Aberrant expression of snoRNAs has also been associated with the development of different diseases and a recent study found alterations in the snoRNA profile of OA joints in mice when compared to healthy controls, highlighting the potential of snoRNAs to be used as novel markers for this disease [21].

Equine miRNAs have been identified in numerous healthy tissues [22, 23] and their potential role in different diseases such as osteochondrosis, rhabdomyolysis and insulin resistance has also been investigated [24–26]. However, information on miRNA influence on the pathogenesis of equine OA is still lacking. Synovial fluid represents a reliable source of chemical information that can accurately reflect pathological conditions affecting the joint due to its functional proximity within joint tissues [27]. In 2010, Murata et al. investigated the presence and stability of miRNAs in synovial fluid for the first time, and found five differentially expressed miRNAs in human OA patients compared to healthy controls, supporting the potential use of synovial fluid miRNAs as diagnostic biomarkers [11]. More recently, a screening of 752 miRNAs in synovial fluid from human patients with early- and late-stage OA demonstrated seven upregulated miRNAs in late-stage OA, irrespective of age, gender and body mass index [28]. Intra-articular treatment with hyaluronic acid was shown to modified miRNA expression in OA patients [29]. Although miRNA expression has not yet been investigated in equine OA, a preliminary study has recently described a reproducible method for miRNA isolation from equine synovial fluid and blood plasma [30].

With growing evidence of alterations in small non-coding RNA patterns in the synovial fluid of OA joints, we theorised that early stages of OA would affect these molecules and potentially provide early biomarkers for OA. Examining expression of small non-coding RNAs in synovial fluid in early OA may also provide further insights on the pathological changes that occur. Therefore, we investigated the profile of small non-coding RNAs of early equine OA synovial fluid using next generation sequencing.

## RESULTS

### Macroscopic and histological assessment

The donors used for small RNA sequencing were selected from an elderly population of horses to account for any age-related changes. The ages of the control (mean±standard deviation; 22±2) and early OA (27±7.5) groups were not significantly different. Because OA is an age-related disease, horses included in the control group presented minor macroscopic or histological changes. For samples used for small RNA sequencing there was a significant increase in the macroscopic score between control 1.0±0.5, early OA; 5.4±1.9 (P=0.04) donors. There was a significant increase in histological score between control; 2.1±0.7, early OA; 6.1±1.5, (P=0.01) (Additional File 1). For the independent cohort the ages of the control and early OA groups were not significantly different (Additional File 1). There was a significant increase in the macroscopic score between control 1.75±1.5, and early OA; 3.6±0.9 (P=0.04) donors and in histological score between control; 1.5 ±1.3, and early OA; 5.8±2.5, (P=0.02) (Additional File 1).

### Analysis of small RNA sequencing data

Summaries of raw, trimmed reads and mapped reads to the Equus caballus database are in Additional File 2. There were 323 small non-coding RNAs identified. The categories of RNA identified are in Figure 1A and included small non-coding RNAs; miRNAs, snoRNAs and small nuclear RNAs (snRNAs).

**Figure 1.**
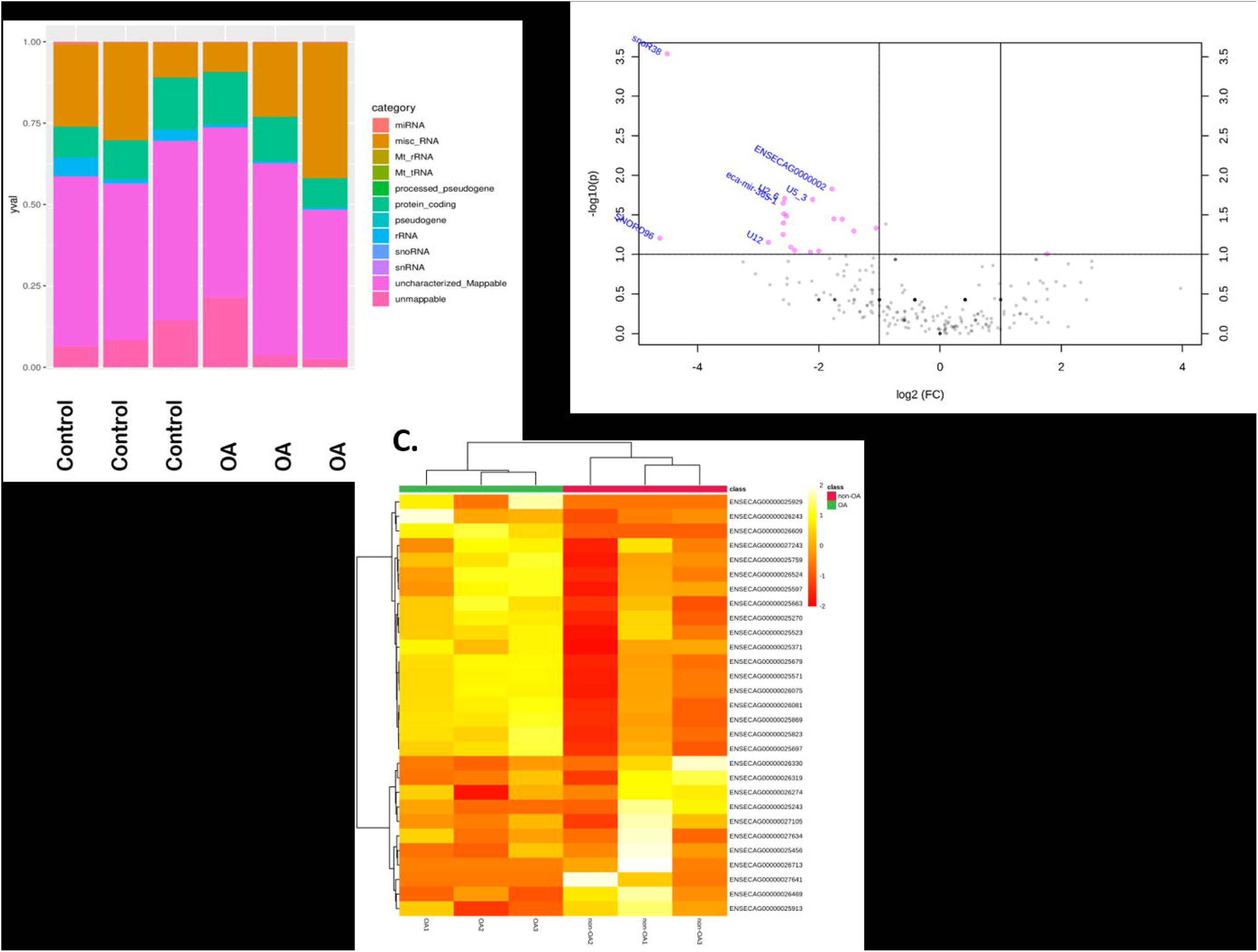
Overview of HiSeq data from equine synovial fluid in control and early OA. (A) Categories of RNAs identified in normal and early OA synovial fluid. (B) Volcano plot of small non-coding RNAs identified represents logFC and –log10 P value. Pink dots represent differentially expressed small non-coding RNAs. (C) A heatmap representation of the differentially expressed small non-coding RNA reads from control (non-OA) and early OA equine synovial fluid. Two-dimensional grid matrix displaying columns referring to the control (non-OA) and early OA samples and rows of small non-coding RNAs identified by their Ensembl identification. The heatmap was generated using log-transformed normalised read counts, normalisation was performed by EdgeR’s trimmed mean of M values. The colour of each entry is determined by the number of reads, ranging from red (negative values) to yellow (positive values).

In total, the expression of 22 small noncoding RNAs; snoRNAs, snRNAs and miRNAs were significantly different in early OA synovial fluid (±1.3 log2 fold change (logFC), and P < 0.05) (Figure 1B; Table 1). We further generated a heatmap of the differentially expressed small non-coding RNAs (Figure 1C).

**Table 1.** Differentially expressed small non-coding RNAs in early OA synovial fluid.

| Ensembl Gene Identification | Gene Name | Gene Biotype | logFC early versus control | P value early versus control |
| --- | --- | --- | --- | --- |
| ENSECAG00000025823 | eca-let-7a-2 | miRNA | 1.39 | 0.02 |
| ENSECAG00000026330 | eca-mir-10a | miRNA | -2.49 | 0.00 |
| ENSECAG00000026319 | eca-mir-125a | miRNA | -1.43 | 0.05 |
| ENSECAG00000026274 | eca-mir-143 | miRNA | -1.87 | 0.04 |
| ENSECAG00000026469 | eca-mir-223 | miRNA | -2.00 | 0.01 |
| ENSECAG00000025270 | eca-mir-23b | miRNA | 1.77 | 0.03 |
| ENSECAG00000025913 | eca-mir-378 | miRNA | -1.34 | 0.04 |
| ENSECAG00000025243 | eca-mir-99a-2 | miRNA | -1.29 | 0.02 |
| ENSECAG00000025456 | ENSECAG00000025456 | miRNA | -1.74 | 0.02 |
| ENSECAG00000025697 | ENSECAG00000025697 | miRNA | 1.31 | 0.03 |
| ENSECAG00000025869 | ENSECAG00000025869 | miRNA | 1.44 | 0.02 |
| ENSECAG00000026713 | ENSECAG00000026713 | miRNA | -7.27 | 0.03 |
| ENSECAG00000027105 | ENSECAG00000027105 | miRNA | -1.34 | 0.04 |
| ENSECAG00000027634 | ENSECAG00000027634 | miRNA | -1.77 | 0.04 |
| ENSECAG00000027641 | SCARNA3 | snoRNA | -7.28 | 0.03 |
| ENSECAG00000026609 | snoR38 | snoRNA | 8.01 | 0.01 |
| ENSECAG00000025929 | SNORD96 | snoRNA | 7.61 | 0.01 |
| ENSECAG00000027243 | snoU13 | snoRNA | 2.02 | 0.04 |
| ENSECAG00000025371 | U11 | snRNA | 1.45 | 0.03 |
| ENSECAG00000025759 | U12 | snRNA | 2.70 | 0.00 |
| ENSECAG00000025571 | U2 | snRNA | 2.57 | 0.00 |
| ENSECAG00000025679 | U2 | snRNA | 2.49 | 0.00 |
| ENSECAG00000026075 | U2 | snRNA | 2.52 | 0.00 |
| ENSECAG00000026524 | U2 | snRNA | 2.43 | 0.00 |
| ENSECAG00000026243 | U2 | snRNA | 2.51 | 0.00 |
| ENSECAG00000025523 | U2 | snRNA | 1.48 | 0.03 |
| ENSECAG00000025597 | U2 | snRNA | 1.57 | 0.04 |
| ENSECAG00000025663 | U5 | snRNA | 3.11 | 0.00 |
| ENSECAG00000026081 | U5 | snRNA | 1.92 | 0.01 |

### Confirmation of differential gene expression using qRT-PCR

Seven small non-coding RNAs (miR-143, miR-223, miR-99a, miR-23b, let-7a-2, snord96A, snord13) were selected for further validation using an independent cohort of synovial fluid samples from control (n=6, histological score 1.5 ±1.3) and early OA (n=6, histological score 5.8±2.5) synovial fluid. In agreement with the sequencing data miR-223 was significantly reduced in early OA and miR-23b, let-7a-2, snord96A and snord13 were significantly increased in early OA (Figure 2). For two miRNAs miR-143 and miR-99a-2 quantitative reverse transcription-polymerase chain reaction (qRT-PCR) findings did not validate sequencing findings.

**Figure 2.**
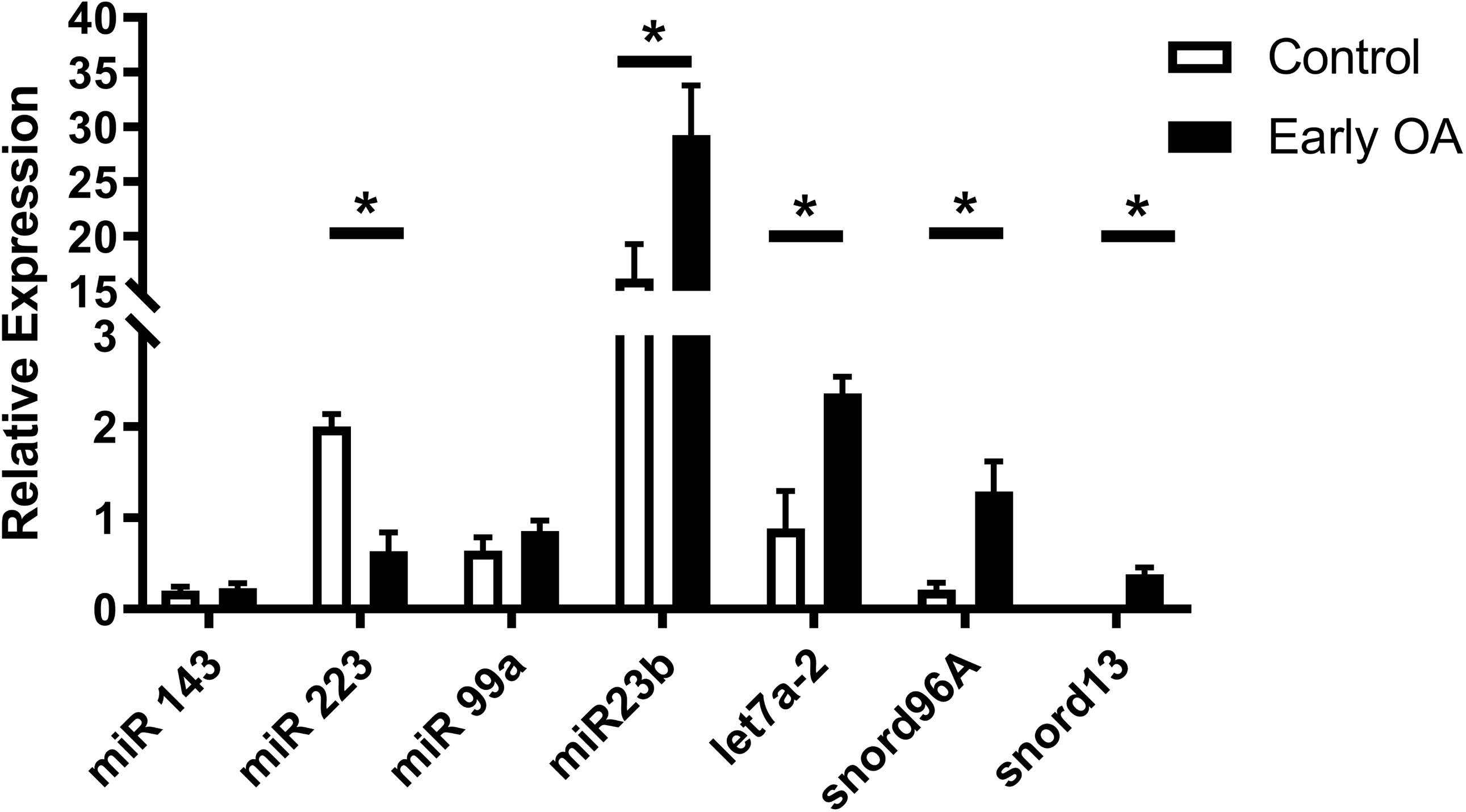
Validation of small non-coding RNAs differentially expressed following small RNA sequencing in an independent cohort using qRT-PCR. RNA extracted from the synovial fluid of six healthy control donors and six early OA donors. Histograms of the relative expression calculated using 2^-DCT method using the geometric mean of miR-100 and miR191 as an endogenous control. All qRT-PCR reactions were performed in triplicate. Statistical significance was tested in Graphpad Prism using a Mann Whitney test. P<0.05; *

### Identification of potential target mRNA genes of the differentially expressed miRNAs

To explore potential biological associations of the differentially expressed miRNAs in early OA synovial fluid we undertook an Ingenuity Pathway Analysis (IPA) ‘Core Analysis’ on these. Interesting features were determined from the gene networks inferred. Significant cellular functions deduced by the differentially expressed miRNAs included apoptosis (P<0.0003), necrosis (P<0.0009), autophagy (P<0.0007) and inflammation (P<0.00001) (Figure 3A).

**Figure 3.**
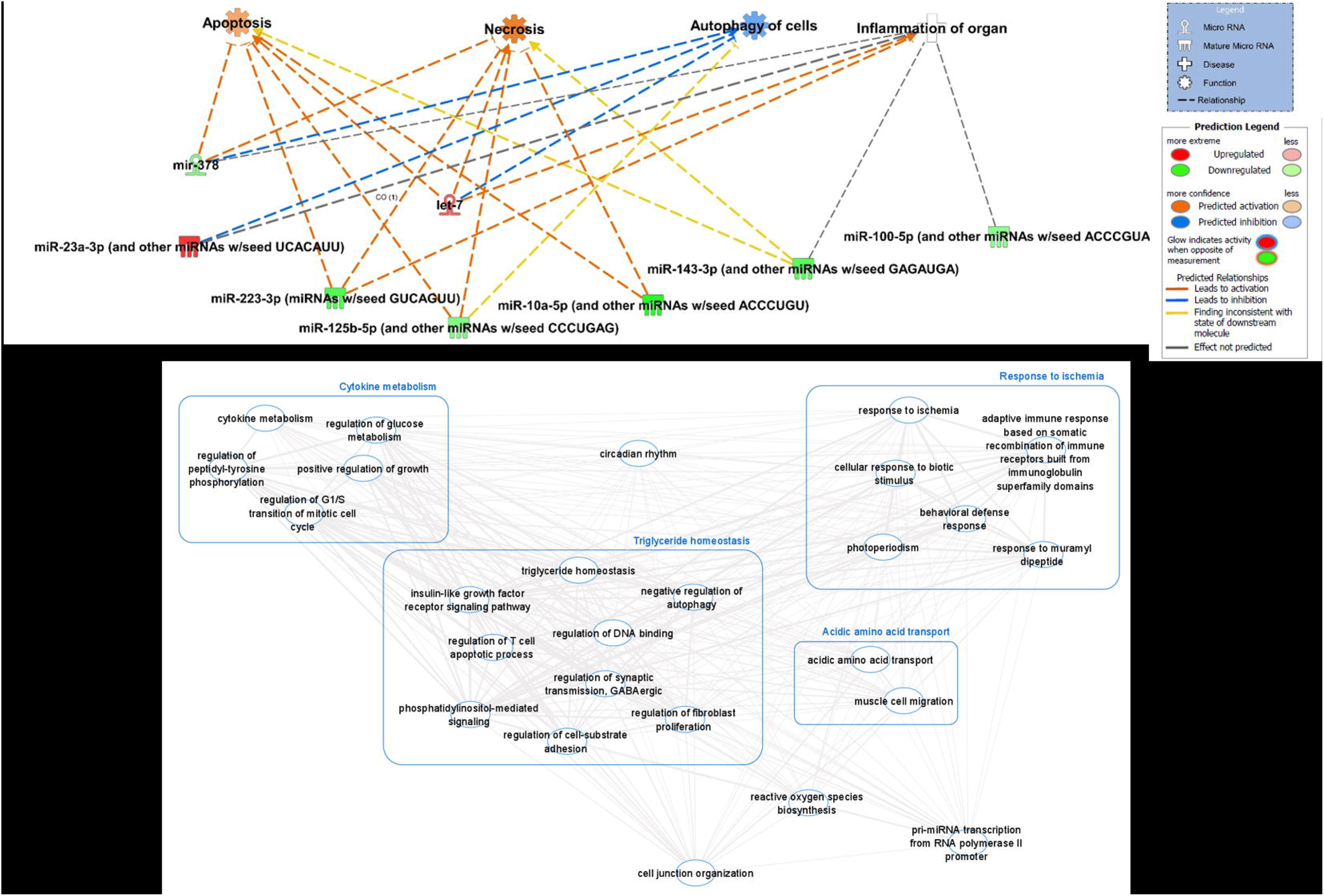
(A) Ingenuity Pathway Analysis (IPA) derived functions of differentially expressed miRNAs in early OA synovial fluid. IPA identified that cellular functions apoptosis, necrosis, autophagy and inflammation were associated with the differentially expressed miRNAs. Figures are graphical representations of molecules identified in our data in their respective networks. Red nodes; upregulated in early OA, and green nodes; downregulated gene expression in early OA synovial fluid. Intensity of colour is related to higher fold-change. Legends to the main features in the networks are shown. The functions colour is dependent on whether it is predicted to be activated or inhibited. (B) The position of differentially expressed miRNAs in the chondrocyte/fibroblast/osteoblast expression network. PANTHER was used to identify gene ontology (GO) biological processes associated with predicted mRNA targets and perform overrepresentation analysis to highlight the GO terms most significantly affected by dysregulated miRNA-mRNA interactions in early OA synovial fluid. GO terms (FDR< 0.05) were summarised and visualised using REViGO and Cytoscape. Allowed similarity setting in REViGO was tiny (0.4). The line width specified the amount of similarity.

Next, we undertook analysis to determine the mRNA targets of the differentially expressed miRNAs. Eight miRNAs were differentially expressed in early OA compared to non-OA controls. Once a conservative filter was applied (only miRNAs with experimentally confirmed or highly conserved predicted targets), six miRNAs remained which collectively putatively target 993 mRNAs. We then additionally added the filters chondrocytes, fibroblast and osteoblasts, removed duplicates and obtained a list of 57 mRNA targets (Additional File 3).

The presumed target mRNAs were input into the gene ontology (GO) tool PANTHER and the biological processes were summarised in REViGO and visualised using Cytoscape (Figure 3B). The top biological processes were regulation of cell population proliferation (false discovery rate (FDR)-adjusted P= 6.24E^-13^), cellular response to chemical stimulus (FDR = 4.54E^-12^) and cell surface receptor signalling pathway (FDR= 6.39E^-12^) (Additional File 4).

## DISCUSSION

The inability to detect pre-clinical changes in OA has been one of the main impediments to the development of effective therapies against this disease [31]. From a biomarker perspective, profiling synovial fluid circulating locally within the affected joint cavity at an early stage may provide new insights into pathological changes occurring during OA initiation and progression, and ultimately allow for the implementation of new therapeutic approaches. Our study is, to the best of our knowledge, the first to characterise the small non-coding RNA profile of synovial fluid in early OA in horses, providing evidence of a pattern of differential expressed synovial fluid miRNAs and other small non-coding RNAs in early OA synovial fluid when compared to our control samples.

Due to the considerable interest in miRNA-mediated gene regulation in recent years, the list of miRNAs possibly implicated in OA and other joint related pathologies has grown [13]. miRNAs that are differentially expressed in joint tissues of patients with OA are likely to contribute to OA pathophysiology and may be utilised as diagnostic factors [32]. One example is miR-140, which is significantly downregulated in human OA cartilage [10]; miR- 140 knockdown in mice promoted proteoglycan loss by disrupting A Disintegrin-like and Metalloproteinase with Thrombospondin Type 1 Motif 5 (ADAMTS-5) expression [33]; intra-articular injection of miR-140 in rats attenuated OA progression by modulating extracellular matrix (ECM) homeostasis [34]; pretransfection with miR-140 partially retarded chondrocyte senescence in an interleukin(IL)-1β induced in vitro model of OA [35]; dysregulation of miR- 140-3p and-5p in synovial fluid has been correlated with OA severity [36]; and a miRNA assay on human serum has identified miR-140-3p as a potential biomarker for OA [37].

In our study, we compared two groups of horses (control vs early OA) based on macroscopic and histological scoring of joints in relation to arthritic changes. Because OA is an age-related disease, both groups were selected from an elderly population and age-matched to account for age-related changes; hence classifying the normal group as “control” as opposed to “healthy”.

Among the differentially expressed miRNAs found in our study, miR-23b was significantly increased in the early OA cohort. miR-23b is thought to be involved in OA progression by targeting cartilage-associated protein (CRTAP) and thus influencing cartilage homeostasis [38]. This miRNA has also been shown to positively regulate the chondrogenic differentiation of mesenchymal stem cells by regulating the expression of sex-determining region Y-Box 9 (SOX9) and protein kinase A (PKA) [39, 40].

Likewise, we found let-7a-2 to be upregulated in early OA. In an experiment comparing miRNA expression in synovial fluid from OA patients undergoing hyaluronic acid treatment, let-7a was significantly upregulated in synovial fluid of OA samples compared to healthy controls; levels of let-7a in affected patients returned to normal after hyaluronan injection [29]. Let-7a is thought to regulate IL-6 receptor (IL6R), and its inhibition can enhance cell proliferation, reduce apoptosis and inhibit inflammatory response in ATDC5 cells in a lipopolysaccharide-induced in vitro model of OA [41]. Members of the let-7 family have often been described in studies involving OA; a large population-based study identified serum let-7e as a promising candidate to predict OA risk, independent of age, sex and body mass index [42]. A recent investigation supported this claim, providing further evidence of decreased expression of let-7e in serum of patients affected with knee OA [43]. The exact roles of miRNAs of the let-7 family remain unclear, but the evidence for their use as biomarkers for OA is growing.

miR-223 was also significantly upregulated in synovial fluid of OA patients prior to intra-articular injection of hyaluronan [29]. miR-223 participates in cartilage homeostasis and structure by targeting growth differentiation factor 5 (GDF5) [38]. Early-stage OA patients showed upregulation of miR-223 in peripheral blood mononuclear cells, with its expression decreasing as OA progressed [44]. In our study, we found miR-223 to be downregulated in the synovial fluid of the early OA cohort, which supports the involvement of this miRNA in the early osteoarthritic process. miR-223 is also enriched in the synovial fluid of patients with rheumatoid arthritis when compared to OA, and can differentiate between patient cohorts [11].

We have previously shown the involvement of snoRNAs in cartilage ageing and OA and their potential use as biomarkers for OA [21]. In this study we identified for the first time snord13 and snord96a as highly expressed small non-coding RNAs in early OA. Our previous work in human OA cartilage identified a dysregulation in SNORD96A expression in ageing and OA. In addition, we demonstrated changes in chondrogenic, hypertrophic, rRNA and OA related gene expression following overexpression and knockdown of SNORD96A in human chondrocytes. Interestingly we also identified an increase in SNORD96A in chondrocytes treated with OA synovial fluid [45]. In another microarray study of young compared to old OA cartilage we identified SNORD13 was increased in OA cartilage [46]. Together these findings indicate that changes in synovial fluid snoRNAs could in part be due to a dysregulation in their expression in cartilage in OA. snoRNAs are emerging with unappreciated functional roles in cell physiology [47] and our results support our earlier work for the potential use of snoRNAs as novel biomarkers in OA [21].

Profiling circulating, cell-free small non-coding RNAs is generally a challenging task due to the limited amount of RNA present in biofluids, as well as presence of inhibitory compounds which potentially hinder downstream enzymatic processes. However, liquid biopsies have gained prominence due to their ease of collection and potential use as diagnostic tools. The horse is a well-established model for spontaneous OA and the horse joint is often used to investigate OA pathogenesis and therapeutics [7]. The metacarpophalangeal joint’s relatively simple structure is ideal for the investigation of early joint modifications because we are unable to determine which tissues are contributing to the overall expression of small non-coding RNAs. This allowed us to more easily determine the origin of small non-coding RNAs that are specifically implicated in OA to cartilage, subchondral bone and synovium.

Future studies would benefit from analysing larger cohorts of patients; our study was limited by the availability of joints with early OA, resulting in a small sample size. Nevertheless, it enabled us to identify small non-coding RNA changes in the initial and an additional cohort and revealed, for the first time, the potential use of small non-coding RNAs as biomarkers for early OA.

Predicted targets of the miRNAs of interest appear to be involved in processes of cellular destruction and inflammation; comparable processes have been previously shown to contribute to the pathogenesis of OA such as degeneration of ECM promoted through pro-inflammatory cytokines [48], synovial inflammation [49] and chondrocyte apoptosis [50], among others. Experimental validation of the predicted target genes can clarify biological mechanisms behind these small non-coding RNAs and elucidate their role in the pathogenesis of OA, which is critical for the success of future interventions, as these molecules can be targeted in a specific manner [51, 52].

These results support the use of synovial fluid small non-coding RNAs as molecular biomarkers for early disease in OA joints. Our future research is currently ascertaining the applicability of these findings in a clinical setting.

## CONCLUSIONS

This study demonstrates that equine synovial fluid displays a pattern of small non-coding RNA differential expression in early OA when compared to controls, as defined by gross and histological scoring and many of these small non-coding RNAs have previously been demonstrated to have a role in OA. The affected biological cellular processes in response to changing miRNAs and their target genes might play an important role in early OA pathogenesis. This opens the possibility of a relatively non-invasive method for early detection of OA. Furthermore, characterisation of these dynamic molecular changes could provide novel insights on the process and mechanism of early OA development.

## METHODS

All reagents were from ThermoFisher Scientific, unless stated.

### Sample collection and preparation

Samples were collected from the metacarpophalangeal joints of horses from an abattoir as a by-product of the agricultural industry. Specifically, the Animal (Scientific procedures) Act 1986, Schedule 2, does not define collection from these sources as scientific procedures. Ethical approval was therefore not required.

Synovial fluid was collected from the metacarpophalangeal joints of control (non-OA), n= 3 (age mean± standard deviation; 22±2 years) and early OA, n=3 (22±7.5 years) non-Thoroughbred horses. At collection synovial fluid was immediately placed on ice and then centrifuged for 10 min at 3000 g and 4°C. The supernatant was removed and stored at - 80°C. Donors were assigned to groups based on scoring of the metacarpophalangeal joint using a macroscopic grading system as previously described [53] and histological scoring system [54].

### RNA isolation, cDNA library preparation and small RNA sequencing

Synovial fluid was treated to reduce viscosity with 1µg/ml of hyaluronidase at 37°C for 1 hr, centrifuged at 1000 g for 5 min, and supernatant used for total RNA extraction using miRNeasy serum kits (Qiagen, Crawley, UK). The integrity of the RNA was assessed on the Agilent 2100 Bioanalyzer system using an RNA Pico chip. 100ng samples were submitted for library preparation using NEBNext® Small RNA Library Prep Set for Illumina (New England Biosciences (NEB), Ipswich, USA) but with the addition of a Cap-Clip™ Acid Pyrophosphatase (Cell script, Madison, USA) step to remove any 5’ cap structures [21] and size selected using a range 120-300bp. This enabled both miRNAs and snoRNAs to be identified in a non-biased approach. The pooled libraries were sequenced on an Illumina HiSeq4000 platform with version 1 chemistry to generate 2 × 150 bp paired-end reads. Data has been submitted to National Centre for Biotechnology Information; accession E-MTAB-8409.

### Small RNA sequencing data analysis

Sequence data were processed through a number of steps to obtain non-coding RNA expression values including; basecalling and de-multiplexing of indexed reads using CASAVA version 1.8.2; adapter and quality trimming using Cutadapt version 1.2.1 [55] and Sickle version 1.200 to obtain fastq files of trimmed reads; aligning reads to horse genome reference sequences (release 90) from Ensembl using Tophat version 2.0.10 [56] with option “–g 1”; counting aligned reads using HTSeq-count [57] against the features defined in horse genome GTF file (release 90).

Differential expression analysis was performed in R using package DESeq2 [58]. The processes and technical details of the analysis include; assessing data variation and detecting outlier samples through comparing variations of within and between sample groups using principle component analysis (PCA) and correlation analysis; handling library size variation using DESeq2 default method; formulating data variation using negative binomial distributions; modelling data using a generalised linear model; computing logFC values for control versus early OA based on model fitting results through contrast fitting approach, evaluating the significance of estimated logFC values by Wald test; adjusting the effects of multiple tests using FDR approach [59] to obtain FDR adjusted P-values.

The Ensembl horse genome GTF file release 90 does not have mature miRNA features. We linked the defined miRNA primary transcripts to miRBase horse miRNA GFF3 file by feature’s genome coordinates so as to obtain the corresponding mature miRNA.

### qRT-PCR validation

Validation of the selected small RNA sequencing results in an independent cohort of equine metacarpophalangeal synovial fluid was undertaken using qRT-PCR. Six control (non-OA), mean±standard deviation (20.2±2.4 years) and six early OA (20.8±4.1) with macroscopically and histologically graded sample scores similar to those used for sequencing were used. Total RNA was extracted as above. Small non-coding RNAs were chosen based on our current work, level of differential expression and following a literature review of differentially expressed genes. These were miR-143, miR-223, miR-99a, miR-23b, let-7a-2, snord96A and snord13. Primer sequences/assays used can be found in Additional File 5. PolyA cDNA was synthesized using 200ng RNA and the miScript II RT Kit. A mastermix was prepared using the miScript SYBR Green PCR Kit (Qiagen, Crawley, UK) and the appropriate bespoke designed miScript Primer Assays (Qiagen, Crawley, UK). Real-time PCR was undertaken using a LightCycler® 96 system (Roche). Steady-state transcript abundance of potential endogenous control genes was measured in the small RNA sequencing data. Assays for four genes – miR-181a, miR-100, miR-191a and U6 were selected as potential reference genes because their expression was unaltered in this study. Stability of this panel of genes was assessed by applying a gene stability tool RefFinder [60]. The geometric mean of miR-100 and miR-191a was selected as the stable endogenous control. miR-100 has been previously used as a normaliser in a similar study as it was identified by NormFinder as the most stable [28]. Relative expression levels were normalised to the geometric mean of miR- 100 and miR-191 and calculated using the 2^-DCT method [61].

### miRNA target prediction and pathway analysis

Potential biological associations of the differentially expressed miRNAs in early OA synovial fluid were identified using Ingenuity Pathway Analysis (IPA) (IPA, Qiagen Redwood City, CA, USA) ‘Core Analysis’. Canonical pathways, networks, and common upstream regulators were then queried. Additionally in order to identify putative miRNA targets, bioinformatic analysis was performed by uploading differentially expressed miRNA data into the MicroRNA Target Filter module within IPA software This identifies experimentally validated miRNA-mRNA interactions from TarBase, miRecords, and the peer-reviewed biomedical literature, as well as predicted miRNA-mRNA interactions from TargetScan. We used a conservative filter at this point, using only experimentally validated and highly conserved predicted mRNA targets for each miRNA. Targets were then also filtered on the cells chondrocyte, osteoblasts and fibroblasts (the latter two settings were the nearest to bone and synovial cells available for selection), to represent joint cells in contact with synovial fluid. PANTHER (GO Ontology database 2020-02-21) [62] was used for overrepresentation analysis of the mRNA targets using Fisher’s Exact test with FDR correction. This tests whether the input mRNAs associate significantly with specific pathways and generates a list of biological process GO terms. Terms with FDR adjusted P < 0.05 were summarised using REViGO [63] with allowed similarity of 0.4 and visualised using Cytoscape [64].

### Statistical analysis

The heatmap and volcano plots were made using MetaboAnalyst 3.5 (http://www.metaboanalyst.ca) which uses the R package of statistical computing software.30 [65]. For statistical evaluation of gene expression data, following normality testing, Mann-Whitney tests were performed using GraphPad Prism version 8.0 for Windows (GraphPad Software, La Jolla California USA, www.graphpad.com); P values are indicated.

## Supporting information

Summary of raw, trimmed reads and mapped reads

mRNA targets predicted by IPA

PANTHER GO terms FDR-adjusted P < 0.05

Primer sequences/assays used

## LIST OF ABBREVIATIONS

ADAMTS5: A Disintegrin-like and Metalloproteinase with Thrombospondin Type 1 Motif 5
CRTAP: cartilage associated protein
DNA: deoxyribonucleic acid
ECM: extracellular matrix
FDR: false discovery rate
GDF5: growth differentiation factor 5
GO: gene ontology
IPA: Ingenuity Pathway Analysis
IL: interleukin
IL6R: interleukin 6 receptor
logFC: log2 fold change
miRNAs: micro RNAs
OA: osteoarthritis
PKA: protein kinase A
qRT-PCR: quantitative reverse transcription polymerase chain reaction
RNA: ribonucleic acid
sncRNAs: small non-coding RNAs
snoRNAs: small nucleolar RNAs
snRNAs: small nuclear RNAs
SOX9: sex-determining region Y – box 9

## DECLARATIONS

### Ethics approval and consent to participate

Synovial fluid was collected as a by-product of the agricultural industry. Specifically, the Animal (Scientific procedures) Act 1986, Schedule 2, does not define collection from these sources as scientific procedures. Ethical approval was therefore not required.

### Consent for publication

Not applicable.

### Availability of data and materials

Data has been submitted to National Centre for Biotechnology Information; accession E-MTAB-8409.The datasets supporting the conclusions of this article are included within the article and its additional files.

### Authors’ contributions

MP, PC and TW designed and coordinated the study. MP, KB, PD collected the samples. PB, PD and KB processed the samples for small RNA sequencing. PD, CAF, YA and CC processed the samples for validation and performed qRT-PCR. MP and CC conducted the statistical analysis and drafted the manuscript. All authors revised the draft critically and read and approved the final submitted manuscript.

## Acknowledgments

The authors would like to thank staff at both the F Drury and Sons abattoir, Swindon for assistance in sample collection and processing, Valerie Tilston for preparation of histology slides, and Aibek Smagul for bioinformatics support.

## ADDITIONAL FILES

**Additional file 1**. Histograms of age, gross score and Modified Mankin’s Score for dependent and independent equine donor cohorts (.tiff). Expressions are means and error bars ± standard error means. Statistical analysis undertaken in GraphPad Prism 8.0 using a Mann Whitney Test. P values *; P <0.05.

**Additional file 2**. Summary of raw, trimmed reads and mapped reads (.xlsx). Summary of raw, trimmed reads and mapped reads to Equus caballus database, from analysis of small RNA sequencing data.

**Additional file 3**. mRNA targets predicted by IPA (.xlsx). List of mRNA targets predicted by bioinformatic analysis with IPA software, using a conservative filter of only experimentally validated and highly conserved predicted mRNA targets for each miRNA. Targets were then also filtered on the cells chondrocyte, osteoblasts and fibroblasts

**Additional file 4**. PANTHER GO terms FDR-adjusted P < 0.05 (.xlsx). List of GO terms with FDR-adjusted P < 0.05, obtained with PANTHER overrepresentation analysis of the mRNA targets using Fisher’s Exact test.

**Additional file 5**. Primer sequences/assays used for detection of small non-coding RNAs through qRT-PCR analysis (.xlsx). For miRNAs and snoRNAs with sequences homologous to human, Qiagen primer assays were used. Remaining miRNA primers were customised using Eurogentec primer design.

## REFERENCES

1. Ireland JL, Clegg PD, Mcgowan CM, Mckane SA, Chandler KJ, Pinchbeck GL. Disease prevalence in geriatric horses in the United Kingdom: Veterinary clinical assessment of 200 cases. Equine Vet J. 2012;44:101–6.

2. Ireland JL, Clegg PD, McGowan CM, Platt L, Pinchbeck GL. Factors associated with mortality of geriatric horses in the United Kingdom. Prev Vet Med. 2011;101:204–18.

3. Woolf AD, Pfleger B. Burden of major musculoskeletal conditions. Bull World Health Organ. 2003;81:646–56.

4. Mobasheri A, Batt M. An update on the pathophysiology of osteoarthritis. Ann Phys Rehabil Med. 2016;59:333–9.

5. Ashkavand Z, Malekinejad H, Vishwanath BS. The pathophysiology of osteoarthritis. J Pharm Res. 2013;7:132–8.

6. Goodrich LR, Nixon AJ. Medical treatment of osteoarthritis in the horse - A review. Vet J. 2006;171:51–69.

7. McIlwraith CW, Frisbie DD, Kawcak CE. The horse as a model of naturally occurring osteoarthritis. Bone Joint Res. 2012;1:297–309.

8. Adams BD, Parsons C, Walker L, Zhang WC, Slack FJ. Targeting noncoding RNAs in disease. J Clin Invest. 2017;127:761–71.

9. Díaz-Prado S, Cicione C, Muiños-López E, Hermida-Gómez T, Oreiro N, Fernández-López C, et al. Characterization of microRNA expression profiles in normal and osteoarthritic human chondrocytes. BMC Musculoskelet Disord. 2012;13:144.

10. Miyaki S, Nakasa T, Otsuki S, Grogan SP, Higashiyama R, Inoue A, et al. MicroRNA-140 is expressed in differentiated human articular chondrocytes and modulates interleukin-1 responses. Arthritis Rheum. 2009;60:2723–30.

11. Murata K, Yoshitomi H, Tanida S, Ishikawa M, Nishitani K, Ito H, et al. Plasma and synovial fluid microRNAs as potential biomarkers of rheumatoid arthritis and osteoarthritis. Arthritis Res Ther. 2010;12:R86.

12. Yu X-M, Meng H-Y, Yuan X-L, Wang Y, Guo Q-Y, Peng J, et al. MicroRNAs’ Involvement in Osteoarthritis and the Prospects for Treatments. Evid Based Complement Alternat Med. 2015;2015:236179.

13. Endisha H, Rockel J, Jurisica I, Kapoor M. The complex landscape of microRNAs in articular cartilage: biology, pathology, and therapeutic targets. JCI insight. 2018;3.

14. Peffers MJ, Balaskas P, Smagul A. Osteoarthritis year in review 2017: genetics and epigenetics. Osteoarthr Cartil. 2018;26:304–11.

15. Ge Q, Zhou Y, Lu J, Bai Y, Xie X, Lu Z. miRNA in Plasma Exosome is Stable under Different Storage Conditions. Molecules. 2014;19:1568–75.

16. Wang K. The Ubiquitous Existence of MicroRNA in Body Fluids. Clin Chem. 2017;63:784–5.

17. Zhang Z, Qin YW, Brewer G, Jing Q. MicroRNA degradation and turnover: Regulating the regulators. Wiley Interdiscip Rev RNA. 2012;3:593–600.

18. Moldovan L, Batte KE, Trgovcich J, Wisler J, Marsh CB, Piper M. Methodological challenges in utilizing miRNAs as circulating biomarkers. J Cell Mol Med. 2014;18:371–90.

19. Buschmann D, Haberberger A, Kirchner B, Spornraft M, Riedmaier I, Schelling G, et al. Toward reliable biomarker signatures in the age of liquid biopsies - How to standardize the small RNA-Seq workflow. Nucleic Acids Res. 2016;44:5995–6018.

20. Stepanov GA, Filippova JA, Komissarov AB, Kuligina E V, Richter VA, Semenov D V. Regulatory Role of Small Nucleolar RNAs in Human Diseases. Biomed Res Int. 2015;2015:206849. doi:10.1155/2015/206849.

21. Steinbusch MMF, Fang Y, Milner PI, Clegg PD, Young DA, Welting TJM, et al. Serum snoRNAs as biomarkers for joint ageing and post traumatic osteoarthritis. Sci Rep. 2017;7:1–11.

22. Kim M-C, Lee S-W, Ryu D-Y, Cui F-J, Bhak J, Kim Y. Identification and Characterization of MicroRNAs in Normal Equine Tissues by Next Generation Sequencing. PLoS One. 2014;9:e93662. doi:10.1371/journal.pone.0093662.

23. Pacholewska A, Mach N, Mata X, Vaiman A, Schibler L, Barrey E, et al. Novel equine tissue miRNAs and breed-related miRNA expressed in serum. BMC Genomics. 2016;17:1–15.

24. Barrey E, Bonnamy B, Barrey EJ, Mata X, Chaffaux S, Guerin G. Muscular microRNA expressions in healthy and myopathic horses suffering from polysaccharide storage myopathy or recurrent exertional rhabdomyolysis. Equine Vet J. 2010;42 SUPPL. 38:303–10.

25. Desjardin C, Vaiman A, Mata X, Legendre R, Laubier J, Kennedy SP, et al. Next-generation sequencing identifies equine cartilage and subchondral bone miRNAs and suggests their involvement in osteochondrosis physiopathology. BMC Genomics. 2014;15:798. doi:10.1186/1471-2164-15-798.

26. da Costa Santos H, Hess T, Bruemmer J, Splan R. Possible Role of MicroRNA in Equine Insulin Resistance: A Pilot Study. J Equine Vet Sci. 2018;63:74–9.

27. McIlwraith CW. Use of synovial fluid and serum biomarkers in equine bone and joint disease: a review. Equine Vet J. 2010;37:473–82.

28. Li Y-H, Tavallaee G, Tokar T, Nakamura A, Sundararajan K, Weston A, et al. Identification of synovial fluid microRNA signature in knee osteoarthritis: differentiating early- and late-stage knee osteoarthritis. Osteoarthr Cartil. 2016;24:1577–86.

29. Xu JF, Zhang SJ, Zhao C, Qiu BS, Gu HF, Hong JF, et al. Altered microRNA Expression Profile in Synovial Fluid from Patients with Knee Osteoarthritis with Treatment of Hyaluronic Acid. Mol Diagnosis Ther. 2015;19:299–308.

30. Antunes J, Koch TG, Koenig J, Cote N, Dubois M-S. On the road to biomarkers: developing a robust system for miRNA evaluation in equine blood and synovial fluid. Osteoarthr Cartil. 2019;27:S110–1.

31. Chu CR, Williams AA, Coyle CH, Bowers ME. Early diagnosis to enable early treatment of pre-osteoarthritis. Arthritis Res Ther. 2012;14:212. doi:10.1186/ar3845.

32. Zhang M, Lygrissea K, Wanga J. Role of MicroRNA in Osteoarthritis. J Arthritis. 2017;06. doi:10.4172/2167-7921.1000239.

33. Miyaki S, Sato T, Inoue A, Otsuki S, Ito Y, Yokoyama S, et al. MicroRNA-140 plays dual roles in both cartilage development and homeostasis. Genes Dev. 2010;24:1173–85.

34. Si HB, Zeng Y, Liu SY, Zhou ZK, Chen YN, Cheng JQ, et al. Intra-articular injection of microRNA-140 (miRNA-140) alleviates osteoarthritis (OA) progression by modulating extracellular matrix (ECM) homeostasis in rats. Osteoarthr Cartil. 2017;25:1698–707.

35. Si H bo, Yang T min, Li L, Tian M, Zhou L, Li D ping, et al. miR-140 Attenuates the Progression of Early-Stage Osteoarthritis by Retarding Chondrocyte Senescence. Mol Ther - Nucleic Acids. 2020;19:15–30.

36. Yin C-M, Suen W-C-W, Lin S, Wu X-M, Li G, Pan X-H. Dysregulation of both miR-140-3p and miR-140-5p in synovial fluid correlate with osteoarthritis severity. Bone Joint Res. 2017;6:612–8.

37. Ntoumou E, Tzetis M, Braoudaki M, Lambrou G, Poulou M, Malizos K, et al. Serum microRNA array analysis identifies miR-140-3p, miR-33b-3p and miR-671-3p as potential osteoarthritis biomarkers involved in metabolic processes. Clin Epigenetics. 2017;9:127.

38. Iliopoulos D, Malizos KN, Oikonomou P, Tsezou A. Integrative MicroRNA and Proteomic Approaches Identify Novel Osteoarthritis Genes and Their Collaborative Metabolic and Inflammatory Networks. PLoS One. 2008;3:e3740. doi:10.1371/journal.pone.0003740.

39. Ham O, Song BW, Lee SY, Choi E, Cha MJ, Lee CY, et al. The role of microRNA-23b in the differentiation of MSC into chondrocyte by targeting protein kinase A signaling. Biomaterials. 2012;33:4500–7.

40. Karlsen TA, Jakobsen RB, Mikkelsen TS, Brinchmann JE. MicroRNA-140 targets RALA and regulates chondrogenic differentiation of human mesenchymal stem cells by translational enhancement of SOX9 and ACAN. Stem Cells Dev. 2014;23:290–304.

41. Sui G, Zhang L, Hu Y. MicroRNA-let-7a inhibition inhibits LPS-induced inflammatory injury of chondrocytes by targeting IL6R. Mol Med Rep. 2019;20:2633–40.

42. Beyer C, Zampetaki A, Lin NY, Kleyer A, Perricone C, Iagnocco A, et al. Signature of circulating microRNAs in osteoarthritis. Ann Rheum Dis. 2015;74:e18–e18.

43. Feng L, Feng C, Wang CX, Xu DY, Chen JJ, Huang JF, et al. Circulating microRNA let–7e is decreased in knee osteoarthritis, accompanied by elevated apoptosis and reduced autophagy. Int J Mol Med. 2020;45:1464–76.

44. Okuhara A, Nakasa T, Shibuya H, Niimoto T, Adachi N, Deie M, et al. Changes in microRNA expression in peripheral mononuclear cells according to the progression of osteoarthritis. Mod Rheumatol. 2012;22:446–57.

45. Peffers MJ, Chabronova A, Balaskas P, Fang Y, Dyer P, Cremers A, et al. SnoRNA signatures in cartilage ageing and osteoarthritis. Preprint at https://www.biorxiv.org/content/10.1101/2020.04.01.019505v1 (2020).

46. Peffers MJ, Ripmeester E, Caron M, Steinbusch M, Balaskas P, Cremers A, et al. A role for the snoRNA U3 in the altered translational capacity of ageing and osteoarthritic chondrocytes. Osteoarthr Cartil. 2018;26:S45–6.

47. Mcmahon M, Contreras A, Ruggero D. Small RNAs with big implications: New insights into H/ACA snoRNA function and their role in human disease. Wiley Interdiscip Rev RNA. 2015;6:173–89.

48. Goldring SR, Goldring MB. The role of cytokines in cartilage matrix degeneration in osteoarthritis. In: Clinical Orthopaedics and Related Research. Lippincott Williams and Wilkins; 2004. p. S27–36.

49. Wang X, Hunter DJ, Jin X, Ding C. The importance of synovial inflammation in osteoarthritis: current evidence from imaging assessments and clinical trials. Osteoarthr Cartil. 2018;26:165–74.

50. Hwang HS, Kim HA. Chondrocyte apoptosis in the pathogenesis of osteoarthritis. Int J Mol Sci. 2015;16:26035–54.

51. Nakamura A, Rampersaud YR, Nakamura S, Sharma A, Zeng F, Rossomacha E, et al. MicroRNA-181a-5p antisense oligonucleotides attenuate osteoarthritis in facet and knee joints. Ann Rheum Dis. 2019;78:111–21.

52. Baek D, Lee KM, Park KW, Suh JW, Choi SM, Park KH, et al. Inhibition of miR-449a Promotes Cartilage Regeneration and Prevents Progression of Osteoarthritis in In Vivo Rat Models. Mol Ther - Nucleic Acids. 2018;13:322–33.

53. Kawcak CE, Frisbie DD, Werpy NM, Park RD, Mcilwraith CW. Effects of exercise vs experimental osteoarthritis on imaging outcomes. Osteoarthr Cartil. 2008;16:1519–25.

54. Mcilwraith CW, Frisbie DD, Kawcak CE, Fuller CJ, Hurtig M, Cruz A. The OARSI histopathology initiative e recommendations for histological assessments of osteoarthritis in the horse. Osteoarthr Cartil. 2010;18:S93–105.

55. Martin M. Cutadapt removes adapter sequences from high-throughput sequencing reads. EMBnet.journal. 2011;17:10. doi:10.14806/ej.17.1.200.

56. Kim D, Pertea G, Trapnell C, Pimentel H, Kelley R, Salzberg SL. TopHat2: accurate alignment of transcriptomes in the presence of insertions, deletions and gene fusions. Genome Biol. 2013;14. doi:10.1186/gb-2013-14-4-r36.

57. Anders S, Pyl PT, Huber W. HTSeq-a Python framework to work with high-throughput sequencing data. Bioinformatics. 2015;31:166–9.

58. Anders S, Huber W. Differential expression analysis for sequence count data. Genome Biol. 2010;11. doi:10.1186/gb-2010-11-10-r106.

59. Benjamini Y, Hochberg Y. Controlling the False Discovery Rate: A Practical and Powerful Approach to Multiple Testing. J R Stat Soc Ser B. 1995;57:289–300.

60. Xie F, Xiao P, Chen D, Xu L, Zhang B. miRDeepFinder: A miRNA analysis tool for deep sequencing of plant small RNAs. Plant Mol Biol. 2012;80:75–84.

61. Livak KJ, Schmittgen TD. Analysis of relative gene expression data using real-time quantitative PCR and the 2-ΔΔCT method. Methods. 2001;25:402–8.

62. Mi H, Muruganujan A, Ebert D, Huang X, Thomas PD. PANTHER version 14: more genomes, a new PANTHER GO-slim and improvements in enrichment analysis tools. Nucleic Acids Res. 2018;47:D419–26.

63. Supek F, Bošnjak M, Škunca N, Šmuc T. REVIGO Summarizes and Visualizes Long Lists of Gene Ontology Terms. PLoS One. 2011;6:e21800. https://doi.org/10.1371/journal.pone.0021800.

64. Shannon P, Markiel A, Ozier O, Baliga NS, Wang JT, Ramage D, et al. Cytoscape: A software Environment for integrated models of biomolecular interaction networks. Genome Res. 2003;13:2498–504.

65. Xia J, Psychogios N, Young N, Wishart DS. MetaboAnalyst: a web server for metabolomic data analysis and interpretation. Nucleic Acids Res. 2009;37. doi:10.1093/nar/gkp356.

