## Supplementary figures and images for "Equine synovial fluid small non-coding RNA signatures in early osteoarthritis"

### Supplemental Data 1

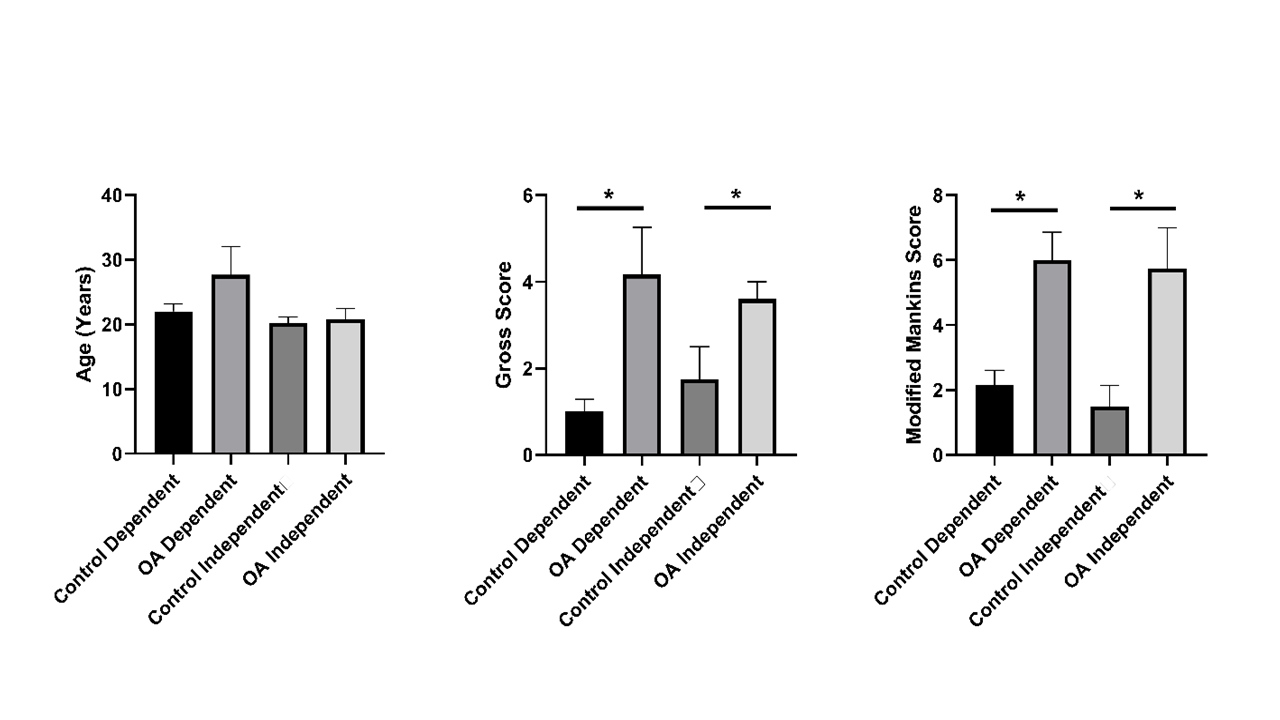
