## Supplementary material for "Equine synovial fluid small non-coding RNA signatures in early osteoarthritis": Summary of raw, trimmed reads and mapped reads

Additional File 2.

| Summaries of raw, trimmed reads and mapped reads | | |  |
| --- | --- | --- | --- |
| Sample | Number of Reads | Number of Reads | Number of Mapped Reads |
| Sample_1-32AB | 140620022 | 123939250 | 56691035 |
| Sample_3-45AB | 127391080 | 105518060 | 37914349 |
| Sample_5-50AB | 144455532 | 123742467 | 58659642 |
| Sample_8-35AB | 198048336 | 184670541 | 88947634 |
| Sample_18-36AB | 155322194 | 142184894 | 63068530 |
| Sample_1016-40AB | 161515408 | 142727461 | 63826484 |
