## Supplementary material for "Equine synovial fluid small non-coding RNA signatures in early osteoarthritis": mRNA targets predicted by IPA

ACVR2A
ADAMTS1
APLN
ARG2
ATP7A
B3GALT4
BCL2
BDNF
BMPR1B
CCR5
CDKN2A
CNR2
CXCL12
CYP1A1
DIO3
ENTPD1
ERBB2
FGFR3
FOSL2
FZD8
GAD2
GJC1
HOXD10
ID2
IGF1R
IGFBP3
IGFBP5
IL18
IL6R
IL6ST
IRS1
KLF4
KRAS
M6PR
MRC1
MTOR
MYD88
MYH1
NCOR2
NF1
NFE2L3
NFIA
NOTCH1
NTRK3
PDE7A
PLAU
PPP3CA
RORA
SDC1
SMAD4

SMO
SP3
SYVN1
TNFAIP6
TP53
TSC22D3
VIM
