## Supplementary material for "Equine synovial fluid small non-coding RNA signatures in early osteoarthritis": PANTHER GO terms FDR-adjusted P < 0.05

Additional File 4

| **Analysis Type:** | **PANTHER Overrepresentation Test (Released 20200407)** | | | | | | | |
|---|---|---|---|---|---|---|---|---|
| **Annotation Version and Release Date:** | **GO Ontology database Released 2020-02-21** |  |  |  |  |  |  |  |
| **Analyzed List:** | **upload_1 (Equus caballus)** | | | | | | | |
| **Reference List:** | **Equus caballus (all genes in database)** | | | | | | | |
| **Test Type:** | **FISHER** | | | | | | | |
| **Correction:** | **FDR** | | | | | | | |
| **GO biological process complete** | | **Equus caballus - REFLIST (21355)** | **upload_1 (56)** | **upload_1 (expected)** | **upload_1 (over/under)** | **upload_1 (fold Enrichment)** | **upload_1 (raw P-value)** | **upload_1 (FDR)** |
| positive regulation of transcription from RNA polymerase II promoter in response to hypoxia | GO:0061419 | 3 | 2 | 0.01 | + | > 100 | 6.69E-05 | 2.94E-03 |
| regulation of cell proliferation involved in heart valve morphogenesis | GO:0003250 | 3 | 2 | 0.01 | + | > 100 | 6.69E-05 | 2.93E-03 |
| negative regulation of neuroblast proliferation | GO:0007406 | 6 | 3 | 0.02 | + | > 100 | 1.41E-06 | 1.20E-04 |
| B cell lineage commitment | GO:0002326 | 4 | 2 | 0.01 | + | > 100 | 1.00E-04 | 4.12E-03 |
| glutamate secretion | GO:0014047 | 4 | 2 | 0.01 | + | > 100 | 1.00E-04 | 4.11E-03 |

| | | | | | | | | |
|---|---|---|---|---|---|---|---|---|
| regulation of retinal cell programmed cell death | GO:0046668 | 4 | 2 | 0.01 | + | > 100 | 1.00E-04 | 4.10E-03 |
| ciliary neurotrophic factor-mediated signaling pathway | GO:0070120 | 5 | 2 | 0.01 | + | > 100 | 1.40E-04 | 5.39E-03 |
| enucleate erythrocyte differentiation | GO:0043353 | 5 | 2 | 0.01 | + | > 100 | 1.40E-04 | 5.37E-03 |
| endocardial cell differentiation | GO:0060956 | 5 | 2 | 0.01 | + | > 100 | 1.40E-04 | 5.36E-03 |
| cardiac endothelial cell differentiation | GO:0003348 | 5 | 2 | 0.01 | + | > 100 | 1.40E-04 | 5.34E-03 |
| atrioventricular valve formation | GO:0003190 | 5 | 2 | 0.01 | + | > 100 | 1.40E-04 | 5.33E-03 |
| positive regulation of astrocyte differentiation | GO:0048711 | 8 | 3 | 0.02 | + | > 100 | 2.76E-06 | 2.22E-04 |
| positive regulation of cell aging | GO:0090343 | 6 | 2 | 0.02 | + | > 100 | 1.86E-04 | 6.77E-03 |
| pri-miRNA transcription by RNA polymerase II | GO:0061614 | 6 | 2 | 0.02 | + | > 100 | 1.86E-04 | 6.75E-03 |
| epithelial to mesenchymal transition involved in endocardial cushion formation | GO:0003198 | 6 | 2 | 0.02 | + | > 100 | 1.86E-04 | 6.73E-03 |
| interleukin-6-mediated signaling pathway | GO:0070102 | 10 | 3 | 0.03 | + | > 100 | 4.76E-06 | 3.35E-04 |

| | | | | | | | | |
|---|---|---|---|---|---|---|---|---|
| negative regulation of production of miRNAs involved in gene silencing by miRNA | GO:1903799 | 10 | 3 | 0.03 | + | > 100 | 4.76E-06 | 3.33E-04 |
| negative regulation of stem cell proliferation | GO:2000647 | 11 | 3 | 0.03 | + | > 100 | 6.04E-06 | 4.11E-04 |
| negative regulation of oligodendrocyte differentiation | GO:0048715 | 11 | 3 | 0.03 | + | > 100 | 6.04E-06 | 4.09E-04 |
| negative regulation of gene silencing by miRNA | GO:0060965 | 12 | 3 | 0.03 | + | 95.33 | 7.54E-06 | 4.87E-04 |
| regulation of chemokine (C-X-C motif) ligand 2 production | GO:2000341 | 8 | 2 | 0.02 | + | 95.33 | 2.98E-04 | 9.88E-03 |
| regulation of transcription from RNA polymerase II promoter in response to hypoxia | GO:0061418 | 8 | 2 | 0.02 | + | 95.33 | 2.98E-04 | 9.86E-03 |
| forebrain morphogenesis | GO:0048853 | 9 | 2 | 0.02 | + | 84.74 | 3.64E-04 | 1.17E-02 |
| regulation of core promoter binding | GO:1904796 | 9 | 2 | 0.02 | + | 84.74 | 3.64E-04 | 1.16E-02 |
| insulin-like growth factor receptor signaling pathway | GO:0048009 | 9 | 2 | 0.02 | + | 84.74 | 3.64E-04 | 1.16E-02 |
| heart valve formation | GO:0003188 | 9 | 2 | 0.02 | + | 84.74 | 3.64E-04 | 1.16E-02 |
| cardiac conduction system development | GO:0003161 | 9 | 2 | 0.02 | + | 84.74 | 3.64E-04 | 1.16E-02 |

| | | | | | | | | |
|---|---|---|---|---|---|---|---|---|
| negative regulation of posttranscriptional gene silencing | GO:0060149 | 14 | 3 | 0.04 | + | 81.72 | 1.12E-05 | 6.73E-04 |
| negative regulation of gene silencing by RNA | GO:0060967 | 14 | 3 | 0.04 | + | 81.72 | 1.12E-05 | 6.70E-04 |
| regulation of astrocyte differentiation | GO:0048710 | 19 | 4 | 0.05 | + | 80.28 | 3.58E-07 | 3.88E-05 |
| natural killer cell differentiation | GO:0001779 | 10 | 2 | 0.03 | + | 76.27 | 4.36E-04 | 1.35E-02 |
| positive regulation of interleukin-17 production | GO:0032740 | 10 | 2 | 0.03 | + | 76.27 | 4.36E-04 | 1.35E-02 |
| left/right axis specification | GO:0070986 | 10 | 2 | 0.03 | + | 76.27 | 4.36E-04 | 1.34E-02 |
| cellular response to interleukin-6 | GO:0071354 | 15 | 3 | 0.04 | + | 76.27 | 1.34E-05 | 7.70E-04 |
| cytokine biosynthetic process | GO:0042089 | 10 | 2 | 0.03 | + | 76.27 | 4.36E-04 | 1.34E-02 |
| endocardial cushion formation | GO:0003272 | 10 | 2 | 0.03 | + | 76.27 | 4.36E-04 | 1.34E-02 |
| endocardium development | GO:0003157 | 10 | 2 | 0.03 | + | 76.27 | 4.36E-04 | 1.34E-02 |
| response to interleukin-6 | GO:0070741 | 16 | 3 | 0.04 | + | 71.5 | 1.59E-05 | 8.88E-04 |

| | | | | | | | | |
|---|---|---|---|---|---|---|---|---|
| negative regulation of neural precursor cell proliferation | GO:2000178 | 16 | 3 | 0.04 | + | 71.5 | 1.59E-05 | 8.85E-04 |
| regulation of neuroblast proliferation | GO:1902692 | 22 | 4 | 0.06 | + | 69.33 | 6.01E-07 | 5.97E-05 |
| epidermis morphogenesis | GO:0048730 | 22 | 4 | 0.06 | + | 69.33 | 6.01E-07 | 5.93E-05 |
| cellular response to sterol | GO:0036315 | 11 | 2 | 0.03 | + | 69.33 | 5.14E-04 | 1.54E-02 |
| cytokine metabolic process | GO:0042107 | 11 | 2 | 0.03 | + | 69.33 | 5.14E-04 | 1.54E-02 |
| astrocyte development | GO:0014002 | 17 | 3 | 0.04 | + | 67.3 | 1.87E-05 | 1.02E-03 |
| regulation of production of miRNAs involved in gene silencing by miRNA | GO:1903798 | 17 | 3 | 0.04 | + | 67.3 | 1.87E-05 | 1.01E-03 |
| regulation of production of small RNA involved in gene silencing by RNA | GO:0070920 | 18 | 3 | 0.05 | + | 63.56 | 2.18E-05 | 1.17E-03 |
| negative regulation of anoikis | GO:2000811 | 12 | 2 | 0.03 | + | 63.56 | 5.99E-04 | 1.76E-02 |
| negative regulation of chemokine production | GO:0032682 | 12 | 2 | 0.03 | + | 63.56 | 5.99E-04 | 1.75E-02 |
| negative regulation of cardiac muscle hypertrophy | GO:0010614 | 12 | 2 | 0.03 | + | 63.56 | 5.99E-04 | 1.75E-02 |

| | | | | | | | | |
|---|---|---|---|---|---|---|---|---|
| positive regulation of smooth muscle cell migration | GO:0014911 | 12 | 2 | 0.03 | + | 63.56 | 5.99E-04 | 1.74E-02 |
| negative regulation of muscle hypertrophy | GO:0014741 | 12 | 2 | 0.03 | + | 63.56 | 5.99E-04 | 1.74E-02 |
| hair follicle morphogenesis | GO:0031069 | 19 | 3 | 0.05 | + | 60.21 | 2.52E-05 | 1.31E-03 |
| regulation of smooth muscle cell migration | GO:0014910 | 26 | 4 | 0.07 | + | 58.67 | 1.09E-06 | 9.59E-05 |
| cardiac left ventricle morphogenesis | GO:0003214 | 13 | 2 | 0.03 | + | 58.67 | 6.90E-04 | 1.96E-02 |
| positive regulation of glial cell differentiation | GO:0045687 | 20 | 3 | 0.05 | + | 57.2 | 2.89E-05 | 1.45E-03 |
| negative regulation of glial cell differentiation | GO:0045686 | 20 | 3 | 0.05 | + | 57.2 | 2.89E-05 | 1.44E-03 |
| regulation of interleukin-17 production | GO:0032660 | 21 | 3 | 0.06 | + | 54.48 | 3.30E-05 | 1.60E-03 |
| regulation of oligodendrocyte differentiation | GO:0048713 | 21 | 3 | 0.06 | + | 54.48 | 3.30E-05 | 1.60E-03 |
| release of cytochrome c from mitochondria | GO:0001836 | 14 | 2 | 0.04 | + | 54.48 | 7.88E-04 | 2.19E-02 |
| regulation of anoikis | GO:2000209 | 14 | 2 | 0.04 | + | 54.48 | 7.88E-04 | 2.19E-02 |

| | | | | | | | | |
|---|---|---|---|---|---|---|---|---|
| regulation of synaptic transmission, GABAergic | GO:0032228 | 14 | 2 | 0.04 | + | 54.48 | 7.88E-04 | 2.19E-02 |
| negative regulation of cell migration involved in sprouting angiogenesis | GO:0090051 | 14 | 2 | 0.04 | + | 54.48 | 7.88E-04 | 2.18E-02 |
| reactive oxygen species biosynthetic process | GO:1903409 | 14 | 2 | 0.04 | + | 54.48 | 7.88E-04 | 2.18E-02 |
| mammary gland epithelial cell differentiation | GO:0060644 | 14 | 2 | 0.04 | + | 54.48 | 7.88E-04 | 2.17E-02 |
| cardiac epithelial to mesenchymal transition | GO:0060317 | 15 | 2 | 0.04 | + | 50.85 | 8.91E-04 | 2.38E-02 |
| circadian behavior | GO:0048512 | 15 | 2 | 0.04 | + | 50.85 | 8.91E-04 | 2.37E-02 |
| negative regulation of gliogenesis | GO:0014014 | 23 | 3 | 0.06 | + | 49.74 | 4.22E-05 | 2.01E-03 |
| negative regulation of gene silencing | GO:0060969 | 24 | 3 | 0.06 | + | 47.67 | 4.74E-05 | 2.23E-03 |
| negative regulation of epidermis development | GO:0045683 | 16 | 2 | 0.04 | + | 47.67 | 1.00E-03 | 2.63E-02 |
| T cell lineage commitment | GO:0002360 | 16 | 2 | 0.04 | + | 47.67 | 1.00E-03 | 2.62E-02 |
| regulation of cell proliferation involved in heart morphogenesis | GO:2000136 | 16 | 2 | 0.04 | + | 47.67 | 1.00E-03 | 2.62E-02 |

| | | | | | | | | |
|---|---|---|---|---|---|---|---|---|
| photoperiodism | GO:0009648 | 16 | 2 | 0.04 | + | 47.67 | 1.00E-03 | 2.61E-02 |
| entrainment of circadian clock by photoperiod | GO:0043153 | 16 | 2 | 0.04 | + | 47.67 | 1.00E-03 | 2.61E-02 |
| rhythmic behavior | GO:0007622 | 16 | 2 | 0.04 | + | 47.67 | 1.00E-03 | 2.60E-02 |
| neurotrophin signaling pathway | GO:0038179 | 17 | 2 | 0.04 | + | 44.86 | 1.12E-03 | 2.86E-02 |
| positive regulation of transcription from RNA polymerase II promoter involved in cellular response to chemical stimulus | GO:1901522 | 17 | 2 | 0.04 | + | 44.86 | 1.12E-03 | 2.85E-02 |
| response to muramyl dipeptide | GO:0032495 | 17 | 2 | 0.04 | + | 44.86 | 1.12E-03 | 2.85E-02 |
| positive regulation of positive chemotaxis | GO:0050927 | 17 | 2 | 0.04 | + | 44.86 | 1.12E-03 | 2.84E-02 |
| activation of protein kinase B activity | GO:0032148 | 17 | 2 | 0.04 | + | 44.86 | 1.12E-03 | 2.84E-02 |
| negative regulation of fibroblast proliferation | GO:0048147 | 17 | 2 | 0.04 | + | 44.86 | 1.12E-03 | 2.83E-02 |
| left/right pattern formation | GO:0060972 | 17 | 2 | 0.04 | + | 44.86 | 1.12E-03 | 2.83E-02 |
| atrioventricular valve morphogenesis | GO:0003181 | 17 | 2 | 0.04 | + | 44.86 | 1.12E-03 | 2.82E-02 |

| | | | | | | | | |
|---|---|---|---|---|---|---|---|---|
| regulation of cardiac muscle hypertrophy | GO:0010611 | 26 | 3 | 0.07 | + | 44 | 5.90E-05 | 2.70E-03 |
| negative regulation of myoblast differentiation | GO:0045662 | 18 | 2 | 0.05 | + | 42.37 | 1.24E-03 | 3.09E-02 |
| regulation of axon regeneration | GO:0048679 | 18 | 2 | 0.05 | + | 42.37 | 1.24E-03 | 3.08E-02 |
| positive regulation of smooth muscle cell proliferation | GO:0048661 | 36 | 4 | 0.09 | + | 42.37 | 3.58E-06 | 2.70E-04 |
| regulation of positive chemotaxis | GO:0050926 | 18 | 2 | 0.05 | + | 42.37 | 1.24E-03 | 3.08E-02 |
| positive regulation of neuron apoptotic process | GO:0043525 | 18 | 2 | 0.05 | + | 42.37 | 1.24E-03 | 3.07E-02 |
| response to sterol | GO:0036314 | 18 | 2 | 0.05 | + | 42.37 | 1.24E-03 | 3.07E-02 |
| muscle cell migration | GO:0014812 | 18 | 2 | 0.05 | + | 42.37 | 1.24E-03 | 3.06E-02 |
| regulation of type 2 immune response | GO:0002828 | 18 | 2 | 0.05 | + | 42.37 | 1.24E-03 | 3.05E-02 |
| homeostasis of number of cells within a tissue | GO:0048873 | 27 | 3 | 0.07 | + | 42.37 | 6.54E-05 | 2.92E-03 |
| regulation of muscle hypertrophy | GO:0014743 | 27 | 3 | 0.07 | + | 42.37 | 6.54E-05 | 2.91E-03 |

| | | | | | | | | |
|---|---|---|---|---|---|---|---|---|
| regulation of hair follicle development | GO:0051797 | 19 | 2 | 0.05 | + | 40.14 | 1.37E-03 | 3.30E-02 |
| cellular response to nerve growth factor stimulus | GO:1990090 | 19 | 2 | 0.05 | + | 40.14 | 1.37E-03 | 3.30E-02 |
| response to nerve growth factor | GO:1990089 | 19 | 2 | 0.05 | + | 40.14 | 1.37E-03 | 3.29E-02 |
| regulation of insulin-like growth factor receptor signaling pathway | GO:0043567 | 19 | 2 | 0.05 | + | 40.14 | 1.37E-03 | 3.29E-02 |
| atrioventricular valve development | GO:0003171 | 19 | 2 | 0.05 | + | 40.14 | 1.37E-03 | 3.28E-02 |
| phosphatidylinositol 3-kinase signaling | GO:0014065 | 29 | 3 | 0.08 | + | 39.45 | 7.97E-05 | 3.41E-03 |
| regulation of glial cell differentiation | GO:0045685 | 39 | 4 | 0.1 | + | 39.11 | 4.80E-06 | 3.33E-04 |
| positive regulation of gliogenesis | GO:0014015 | 30 | 3 | 0.08 | + | 38.13 | 8.75E-05 | 3.69E-03 |
| smooth muscle tissue development | GO:0048745 | 20 | 2 | 0.05 | + | 38.13 | 1.50E-03 | 3.50E-02 |
| regulation of neuron projection regeneration | GO:0070570 | 20 | 2 | 0.05 | + | 38.13 | 1.50E-03 | 3.50E-02 |
| lens fiber cell differentiation | GO:0070306 | 20 | 2 | 0.05 | + | 38.13 | 1.50E-03 | 3.49E-02 |

| | | | | | | | | |
|---|---|---|---|---|---|---|---|---|
| entrainment of circadian clock | GO:0009649 | 20 | 2 | 0.05 | + | 38.13 | 1.50E-03 | 3.48E-02 |
| regulation of T cell apoptotic process | GO:0070232 | 20 | 2 | 0.05 | + | 38.13 | 1.50E-03 | 3.48E-02 |
| regulation of alpha-beta T cell proliferation | GO:0046640 | 20 | 2 | 0.05 | + | 38.13 | 1.50E-03 | 3.47E-02 |
| response to ischemia | GO:0002931 | 20 | 2 | 0.05 | + | 38.13 | 1.50E-03 | 3.47E-02 |
| regulation of heart morphogenesis | GO:2000826 | 31 | 3 | 0.08 | + | 36.9 | 9.57E-05 | 3.99E-03 |
| astrocyte differentiation | GO:0048708 | 31 | 3 | 0.08 | + | 36.9 | 9.57E-05 | 3.98E-03 |
| regulation of gene silencing by miRNA | GO:0060964 | 31 | 3 | 0.08 | + | 36.9 | 9.57E-05 | 3.97E-03 |
| peripheral nervous system development | GO:0007422 | 52 | 5 | 0.14 | + | 36.67 | 3.85E-07 | 4.14E-05 |
| positive regulation of erythrocyte differentiation | GO:0045648 | 21 | 2 | 0.06 | + | 36.32 | 1.64E-03 | 3.75E-02 |
| myelination in peripheral nervous system | GO:0022011 | 21 | 2 | 0.06 | + | 36.32 | 1.64E-03 | 3.74E-02 |
| peripheral nervous system axon ensheathment | GO:0032292 | 21 | 2 | 0.06 | + | 36.32 | 1.64E-03 | 3.74E-02 |

| | | | | | | | | |
|---|---|---|---|---|---|---|---|---|
| endocardial cushion morphogenesis | GO:0003203 | 21 | 2 | 0.06 | + | 36.32 | 1.64E-03 | 3.73E-02 |
| ventricular septum morphogenesis | GO:0060412 | 32 | 3 | 0.08 | + | 35.75 | 1.05E-04 | 4.24E-03 |
| regulation of posttranscriptional gene silencing | GO:0060147 | 33 | 3 | 0.09 | + | 34.67 | 1.14E-04 | 4.54E-03 |
| gastrulation with mouth forming second | GO:0001702 | 22 | 2 | 0.06 | + | 34.67 | 1.79E-03 | 4.01E-02 |
| positive regulation of kidney development | GO:0090184 | 22 | 2 | 0.06 | + | 34.67 | 1.79E-03 | 4.00E-02 |
| triglyceride homeostasis | GO:0070328 | 22 | 2 | 0.06 | + | 34.67 | 1.79E-03 | 4.00E-02 |
| positive regulation of transcription from RNA polymerase II promoter in response to stress | GO:0036003 | 22 | 2 | 0.06 | + | 34.67 | 1.79E-03 | 3.99E-02 |
| acylglycerol homeostasis | GO:0055090 | 22 | 2 | 0.06 | + | 34.67 | 1.79E-03 | 3.99E-02 |
| acidic amino acid transport | GO:0015800 | 22 | 2 | 0.06 | + | 34.67 | 1.79E-03 | 3.98E-02 |
| negative regulation of macroautophagy | GO:0016242 | 22 | 2 | 0.06 | + | 34.67 | 1.79E-03 | 3.97E-02 |
| regulation of gene silencing by RNA | GO:0060966 | 33 | 3 | 0.09 | + | 34.67 | 1.14E-04 | 4.53E-03 |

| | | | | | | | | |
|---|---|---|---|---|---|---|---|---|
| negative regulation of sprouting angiogenesis | GO:1903671 | 22 | 2 | 0.06 | + | 34.67 | 1.79E-03 | 3.97E-02 |
| digestive tract morphogenesis | GO:0048546 | 34 | 3 | 0.09 | + | 33.65 | 1.24E-04 | 4.86E-03 |
| cardiac septum morphogenesis | GO:0060411 | 57 | 5 | 0.15 | + | 33.45 | 5.89E-07 | 5.93E-05 |
| skeletal muscle fiber development | GO:0048741 | 23 | 2 | 0.06 | + | 33.16 | 1.94E-03 | 4.26E-02 |
| heart trabecula morphogenesis | GO:0061384 | 23 | 2 | 0.06 | + | 33.16 | 1.94E-03 | 4.25E-02 |
| ovarian follicle development | GO:0001541 | 35 | 3 | 0.09 | + | 32.69 | 1.34E-04 | 5.20E-03 |
| negative regulation of animal organ morphogenesis | GO:0110111 | 24 | 2 | 0.06 | + | 31.78 | 2.10E-03 | 4.55E-02 |
| behavioral fear response | GO:0001662 | 24 | 2 | 0.06 | + | 31.78 | 2.10E-03 | 4.54E-02 |
| Schwann cell development | GO:0014044 | 24 | 2 | 0.06 | + | 31.78 | 2.10E-03 | 4.54E-02 |
| cellular senescence | GO:0090398 | 24 | 2 | 0.06 | + | 31.78 | 2.10E-03 | 4.53E-02 |
| lipopolysaccharide-mediated signaling pathway | GO:0031663 | 24 | 2 | 0.06 | + | 31.78 | 2.10E-03 | 4.52E-02 |

| | | | | | | | | |
|---|---|---|---|---|---|---|---|---|
| intrinsic apoptotic signaling pathway in response to endoplasmic reticulum stress | GO:0070059 | 24 | 2 | 0.06 | + | 31.78 | 2.10E-03 | 4.51E-02 |
| extrinsic apoptotic signaling pathway via death domain receptors | GO:0008625 | 24 | 2 | 0.06 | + | 31.78 | 2.10E-03 | 4.51E-02 |
| regulation of stem cell proliferation | GO:0072091 | 49 | 4 | 0.13 | + | 31.13 | 1.12E-05 | 6.72E-04 |
| behavioral defense response | GO:0002209 | 25 | 2 | 0.07 | + | 30.51 | 2.26E-03 | 4.83E-02 |
| negative regulation of striated muscle cell differentiation | GO:0051154 | 25 | 2 | 0.07 | + | 30.51 | 2.26E-03 | 4.82E-02 |
| negative regulation of blood vessel endothelial cell migration | GO:0043537 | 25 | 2 | 0.07 | + | 30.51 | 2.26E-03 | 4.82E-02 |
| epithelial cell differentiation involved in kidney development | GO:0035850 | 25 | 2 | 0.07 | + | 30.51 | 2.26E-03 | 4.81E-02 |
| myotube cell development | GO:0014904 | 25 | 2 | 0.07 | + | 30.51 | 2.26E-03 | 4.80E-02 |
| cardiac atrium morphogenesis | GO:0003209 | 25 | 2 | 0.07 | + | 30.51 | 2.26E-03 | 4.79E-02 |
| regulation of smooth muscle cell proliferation | GO:0048660 | 63 | 5 | 0.17 | + | 30.27 | 9.37E-07 | 8.48E-05 |
| positive regulation of tyrosine phosphorylation of STAT protein | GO:0042531 | 38 | 3 | 0.1 | + | 30.11 | 1.68E-04 | 6.23E-03 |

| | | | | | | | | |
|---|---|---|---|---|---|---|---|---|
| regulation of chemokine production | GO:0032642 | 51 | 4 | 0.13 | + | 29.91 | 1.30E-05 | 7.45E-04 |
| regulation of neural precursor cell proliferation | GO:2000177 | 64 | 5 | 0.17 | + | 29.79 | 1.01E-06 | 9.01E-05 |
| regulation of myoblast differentiation | GO:0045661 | 39 | 3 | 0.1 | + | 29.33 | 1.81E-04 | 6.61E-03 |
| regulation of muscle adaptation | GO:0043502 | 39 | 3 | 0.1 | + | 29.33 | 1.81E-04 | 6.59E-03 |
| metanephros development | GO:0001656 | 65 | 5 | 0.17 | + | 29.33 | 1.08E-06 | 9.57E-05 |
| glial cell development | GO:0021782 | 66 | 5 | 0.17 | + | 28.89 | 1.16E-06 | 1.01E-04 |
| positive regulation of receptor signaling pathway via JAK-STAT | GO:0046427 | 53 | 4 | 0.14 | + | 28.78 | 1.50E-05 | 8.43E-04 |
| natural killer cell activation | GO:0030101 | 40 | 3 | 0.1 | + | 28.6 | 1.94E-04 | 6.98E-03 |
| ventricular cardiac muscle tissue development | GO:0003229 | 40 | 3 | 0.1 | + | 28.6 | 1.94E-04 | 6.97E-03 |
| positive regulation of receptor signaling pathway via STAT | GO:1904894 | 54 | 4 | 0.14 | + | 28.25 | 1.60E-05 | 8.86E-04 |
| negative regulation of G1/S transition of mitotic cell cycle | GO:2000134 | 42 | 3 | 0.11 | + | 27.24 | 2.22E-04 | 7.76E-03 |

| | | | | | | | | |
|---|---|---|---|---|---|---|---|---|
| cellular response to hypoxia | GO:0071456 | 58 | 4 | 0.15 | + | 26.3 | 2.09E-05 | 1.13E-03 |
| regulation of gliogenesis | GO:0014013 | 58 | 4 | 0.15 | + | 26.3 | 2.09E-05 | 1.13E-03 |
| cell aging | GO:0007569 | 44 | 3 | 0.12 | + | 26 | 2.53E-04 | 8.69E-03 |
| negative regulation of cell cycle G1/S phase transition | GO:1902807 | 44 | 3 | 0.12 | + | 26 | 2.53E-04 | 8.67E-03 |
| hair follicle development | GO:0001942 | 59 | 4 | 0.15 | + | 25.85 | 2.23E-05 | 1.19E-03 |
| neuron fate commitment | GO:0048663 | 59 | 4 | 0.15 | + | 25.85 | 2.23E-05 | 1.19E-03 |
| cellular response to decreased oxygen levels | GO:0036294 | 59 | 4 | 0.15 | + | 25.85 | 2.23E-05 | 1.18E-03 |
| phosphatidylinositol-mediated signaling | GO:0048015 | 59 | 4 | 0.15 | + | 25.85 | 2.23E-05 | 1.18E-03 |
| regulation of tissue remodeling | GO:0034103 | 45 | 3 | 0.12 | + | 25.42 | 2.70E-04 | 9.10E-03 |
| endothelial cell differentiation | GO:0045446 | 45 | 3 | 0.12 | + | 25.42 | 2.70E-04 | 9.08E-03 |
| negative regulation of autophagy | GO:0010507 | 45 | 3 | 0.12 | + | 25.42 | 2.70E-04 | 9.06E-03 |

| | | | | | | | | |
|---|---|---|---|---|---|---|---|---|
| skin epidermis development | GO:0098773 | 61 | 4 | 0.16 | + | 25.01 | 2.52E-05 | 1.31E-03 |
| inositol lipid-mediated signaling | GO:0048017 | 61 | 4 | 0.16 | + | 25.01 | 2.52E-05 | 1.30E-03 |
| hair cycle process | GO:0022405 | 62 | 4 | 0.16 | + | 24.6 | 2.68E-05 | 1.37E-03 |
| molting cycle process | GO:0022404 | 62 | 4 | 0.16 | + | 24.6 | 2.68E-05 | 1.37E-03 |
| positive regulation of osteoblast differentiation | GO:0045669 | 47 | 3 | 0.12 | + | 24.34 | 3.04E-04 | 1.00E-02 |
| female gonad development | GO:0008585 | 63 | 4 | 0.17 | + | 24.21 | 2.85E-05 | 1.43E-03 |
| cardiac chamber morphogenesis | GO:0003206 | 95 | 6 | 0.25 | + | 24.08 | 2.54E-07 | 2.89E-05 |
| regulation of tyrosine phosphorylation of STAT protein | GO:0042509 | 48 | 3 | 0.13 | + | 23.83 | 3.23E-04 | 1.05E-02 |
| glial cell differentiation | GO:0010001 | 112 | 7 | 0.29 | + | 23.83 | 2.54E-08 | 3.96E-06 |
| regulation of G1/S transition of mitotic cell cycle | GO:2000045 | 81 | 5 | 0.21 | + | 23.54 | 3.02E-06 | 2.36E-04 |
| hair cycle | GO:0042633 | 65 | 4 | 0.17 | + | 23.47 | 3.20E-05 | 1.58E-03 |

| | | | | | | | | |
|---|---|---|---|---|---|---|---|---|
| molting cycle | GO:0042303 | 65 | 4 | 0.17 | + | 23.47 | 3.20E-05 | 1.57E-03 |
| development of primary female sexual characteristics | GO:0046545 | 66 | 4 | 0.17 | + | 23.11 | 3.38E-05 | 1.64E-03 |
| cardiac septum development | GO:0003279 | 84 | 5 | 0.22 | + | 22.7 | 3.58E-06 | 2.69E-04 |
| multicellular organismal response to stress | GO:0033555 | 51 | 3 | 0.13 | + | 22.43 | 3.82E-04 | 1.21E-02 |
| regulation of osteoblast differentiation | GO:0045667 | 86 | 5 | 0.23 | + | 22.17 | 4.00E-06 | 2.91E-04 |
| kidney epithelium development | GO:0072073 | 104 | 6 | 0.27 | + | 22 | 4.22E-07 | 4.51E-05 |
| regulation of fibroblast proliferation | GO:0048145 | 52 | 3 | 0.14 | + | 22 | 4.04E-04 | 1.27E-02 |
| mechanoreceptor differentiation | GO:0042490 | 52 | 3 | 0.14 | + | 22 | 4.04E-04 | 1.26E-02 |
| endothelium development | GO:0003158 | 52 | 3 | 0.14 | + | 22 | 4.04E-04 | 1.26E-02 |
| vasculogenesis | GO:0001570 | 53 | 3 | 0.14 | + | 21.59 | 4.26E-04 | 1.32E-02 |
| response to hypoxia | GO:0001666 | 124 | 7 | 0.33 | + | 21.53 | 4.94E-08 | 7.01E-06 |

| | | | | | | | | |
|---|---|---|---|---|---|---|---|---|
| cellular response to oxygen levels | GO:0071453 | 71 | 4 | 0.19 | + | 21.48 | 4.44E-05 | 2.10E-03 |
| response to decreased oxygen levels | GO:0036293 | 125 | 7 | 0.33 | + | 21.36 | 5.20E-08 | 7.32E-06 |
| ureteric bud development | GO:0001657 | 72 | 4 | 0.19 | + | 21.19 | 4.68E-05 | 2.21E-03 |
| nephron epithelium morphogenesis | GO:0072088 | 54 | 3 | 0.14 | + | 21.19 | 4.49E-04 | 1.37E-02 |
| nephron morphogenesis | GO:0072028 | 54 | 3 | 0.14 | + | 21.19 | 4.49E-04 | 1.37E-02 |
| rhythmic process | GO:0048511 | 126 | 7 | 0.33 | + | 21.19 | 5.48E-08 | 7.49E-06 |
| circadian rhythm | GO:0007623 | 91 | 5 | 0.24 | + | 20.95 | 5.20E-06 | 3.59E-04 |
| mesonephric tubule development | GO:0072164 | 73 | 4 | 0.19 | + | 20.9 | 4.93E-05 | 2.31E-03 |
| mesonephric epithelium development | GO:0072163 | 73 | 4 | 0.19 | + | 20.9 | 4.93E-05 | 2.30E-03 |
| cellular response to interferon-gamma | GO:0071346 | 55 | 3 | 0.14 | + | 20.8 | 4.72E-04 | 1.43E-02 |
| ventricular septum development | GO:0003281 | 55 | 3 | 0.14 | + | 20.8 | 4.72E-04 | 1.43E-02 |

| | | | | | | | | |
|---|---|---|---|---|---|---|---|---|
| regulation of cell cycle G1/S phase transition | GO:1902806 | 92 | 5 | 0.24 | + | 20.72 | 5.47E-06 | 3.75E-04 |
| female sex differentiation | GO:0046660 | 76 | 4 | 0.2 | + | 20.07 | 5.73E-05 | 2.65E-03 |
| positive regulation of phosphatidylinositol 3-kinase signaling | GO:0014068 | 57 | 3 | 0.15 | + | 20.07 | 5.22E-04 | 1.56E-02 |
| negative regulation of ossification | GO:0030279 | 57 | 3 | 0.15 | + | 20.07 | 5.22E-04 | 1.55E-02 |
| mesonephros development | GO:0001823 | 77 | 4 | 0.2 | + | 19.81 | 6.01E-05 | 2.75E-03 |
| nephron epithelium development | GO:0072009 | 77 | 4 | 0.2 | + | 19.81 | 6.01E-05 | 2.74E-03 |
| regulation of gene silencing | GO:0060968 | 58 | 3 | 0.15 | + | 19.72 | 5.48E-04 | 1.62E-02 |
| skeletal muscle tissue development | GO:0007519 | 97 | 5 | 0.25 | + | 19.66 | 7.00E-06 | 4.54E-04 |
| response to oxygen levels | GO:0070482 | 136 | 7 | 0.36 | + | 19.63 | 9.02E-08 | 1.15E-05 |
| liver development | GO:0001889 | 59 | 3 | 0.15 | + | 19.39 | 5.75E-04 | 1.69E-02 |
| metencephalon development | GO:0022037 | 79 | 4 | 0.21 | + | 19.31 | 6.61E-05 | 2.94E-03 |

| | | | | | | | | |
|---|---|---|---|---|---|---|---|---|
| negative regulation of angiogenesis | GO:0016525 | 79 | 4 | 0.21 | + | 19.31 | 6.61E-05 | 2.93E-03 |
| regulation of glucose metabolic process | GO:0010906 | 79 | 4 | 0.21 | + | 19.31 | 6.61E-05 | 2.92E-03 |
| renal system development | GO:0072001 | 199 | 10 | 0.52 | + | 19.16 | 1.52E-10 | 7.22E-08 |
| negative regulation of blood vessel morphogenesis | GO:2000181 | 80 | 4 | 0.21 | + | 19.07 | 6.93E-05 | 3.01E-03 |
| skeletal muscle organ development | GO:0060538 | 101 | 5 | 0.26 | + | 18.88 | 8.45E-06 | 5.36E-04 |
| nephron tubule development | GO:0072080 | 61 | 3 | 0.16 | + | 18.75 | 6.31E-04 | 1.83E-02 |
| axis specification | GO:0009798 | 61 | 3 | 0.16 | + | 18.75 | 6.31E-04 | 1.82E-02 |
| gliogenesis | GO:0042063 | 144 | 7 | 0.38 | + | 18.54 | 1.31E-07 | 1.60E-05 |
| hepaticobiliary system development | GO:0061008 | 62 | 3 | 0.16 | + | 18.45 | 6.60E-04 | 1.89E-02 |
| cardiac chamber development | GO:0003205 | 125 | 6 | 0.33 | + | 18.3 | 1.18E-06 | 1.02E-04 |
| kidney development | GO:0001822 | 189 | 9 | 0.5 | + | 18.16 | 2.15E-09 | 5.00E-07 |

| | | | | | | | | |
|---|---|---|---|---|---|---|---|---|
| regulation of epidermis development | GO:0045682 | 63 | 3 | 0.17 | + | 18.16 | 6.90E-04 | 1.97E-02 |
| canonical Wnt signaling pathway | GO:0060070 | 63 | 3 | 0.17 | + | 18.16 | 6.90E-04 | 1.97E-02 |
| regulation of striated muscle cell differentiation | GO:0051153 | 63 | 3 | 0.17 | + | 18.16 | 6.90E-04 | 1.96E-02 |
| BMP signaling pathway | GO:0030509 | 64 | 3 | 0.17 | + | 17.88 | 7.21E-04 | 2.04E-02 |
| renal tubule development | GO:0061326 | 65 | 3 | 0.17 | + | 17.6 | 7.53E-04 | 2.12E-02 |
| positive regulation of ossification | GO:0045778 | 66 | 3 | 0.17 | + | 17.33 | 7.86E-04 | 2.20E-02 |
| regulation of blood vessel endothelial cell migration | GO:0043535 | 66 | 3 | 0.17 | + | 17.33 | 7.86E-04 | 2.19E-02 |
| negative regulation of vasculature development | GO:1901343 | 89 | 4 | 0.23 | + | 17.14 | 1.03E-04 | 4.20E-03 |
| response to interferon-gamma | GO:0034341 | 67 | 3 | 0.18 | + | 17.07 | 8.19E-04 | 2.23E-02 |
| negative regulation of cell development | GO:0010721 | 203 | 9 | 0.53 | + | 16.91 | 3.91E-09 | 8.54E-07 |
| urogenital system development | GO:0001655 | 226 | 10 | 0.59 | + | 16.87 | 5.00E-10 | 1.61E-07 |

| | | | | | | | | |
|---|---|---|---|---|---|---|---|---|
| mesenchymal cell development | GO:0014031 | 68 | 3 | 0.18 | + | 16.82 | 8.54E-04 | 2.32E-02 |
| negative regulation of leukocyte differentiation | GO:1902106 | 68 | 3 | 0.18 | + | 16.82 | 8.54E-04 | 2.32E-02 |
| epithelial cell proliferation | GO:0050673 | 68 | 3 | 0.18 | + | 16.82 | 8.54E-04 | 2.31E-02 |
| kidney morphogenesis | GO:0060993 | 68 | 3 | 0.18 | + | 16.82 | 8.54E-04 | 2.31E-02 |
| digestive tract development | GO:0048565 | 91 | 4 | 0.24 | + | 16.76 | 1.12E-04 | 4.48E-03 |
| positive regulation of lymphocyte proliferation | GO:0050671 | 92 | 4 | 0.24 | + | 16.58 | 1.17E-04 | 4.61E-03 |
| regulation of circadian rhythm | GO:0042752 | 69 | 3 | 0.18 | + | 16.58 | 8.89E-04 | 2.38E-02 |
| positive regulation of mononuclear cell proliferation | GO:0032946 | 93 | 4 | 0.24 | + | 16.4 | 1.21E-04 | 4.79E-03 |
| regulation of biomineralization | GO:0110149 | 70 | 3 | 0.18 | + | 16.34 | 9.26E-04 | 2.46E-02 |
| regulation of biomineral tissue development | GO:0070167 | 70 | 3 | 0.18 | + | 16.34 | 9.26E-04 | 2.45E-02 |
| cerebellum development | GO:0021549 | 71 | 3 | 0.19 | + | 16.11 | 9.63E-04 | 2.54E-02 |

| | | | | | | | | |
|---|---|---|---|---|---|---|---|---|
| regulation of phosphatidylinositol 3-kinase signaling | GO:0014066 | 71 | 3 | 0.19 | + | 16.11 | 9.63E-04 | 2.54E-02 |
| skin development | GO:0043588 | 142 | 6 | 0.37 | + | 16.11 | 2.41E-06 | 1.99E-04 |
| positive regulation of developmental growth | GO:0048639 | 119 | 5 | 0.31 | + | 16.02 | 1.81E-05 | 9.94E-04 |
| negative regulation of hemopoiesis | GO:1903707 | 96 | 4 | 0.25 | + | 15.89 | 1.37E-04 | 5.27E-03 |
| cell fate commitment | GO:0045165 | 194 | 8 | 0.51 | + | 15.73 | 5.26E-08 | 7.33E-06 |
| digestive system development | GO:0055123 | 97 | 4 | 0.25 | + | 15.73 | 1.42E-04 | 5.38E-03 |
| regulation of cellular carbohydrate metabolic process | GO:0010675 | 97 | 4 | 0.25 | + | 15.73 | 1.42E-04 | 5.36E-03 |
| cellular response to BMP stimulus | GO:0071773 | 73 | 3 | 0.19 | + | 15.67 | 1.04E-03 | 2.69E-02 |
| response to BMP | GO:0071772 | 73 | 3 | 0.19 | + | 15.67 | 1.04E-03 | 2.68E-02 |
| regulation of lymphocyte proliferation | GO:0050670 | 146 | 6 | 0.38 | + | 15.67 | 2.81E-06 | 2.23E-04 |
| somite development | GO:0061053 | 73 | 3 | 0.19 | + | 15.67 | 1.04E-03 | 2.68E-02 |

| | | | | | | | | |
|---|---|---|---|---|---|---|---|---|
| nephron development | GO:0072006 | 98 | 4 | 0.26 | + | 15.56 | 1.47E-04 | 5.53E-03 |
| positive regulation of leukocyte proliferation | GO:0070665 | 98 | 4 | 0.26 | + | 15.56 | 1.47E-04 | 5.51E-03 |
| regulation of mononuclear cell proliferation | GO:0032944 | 148 | 6 | 0.39 | + | 15.46 | 3.03E-06 | 2.35E-04 |
| osteoblast differentiation | GO:0001649 | 74 | 3 | 0.19 | + | 15.46 | 1.08E-03 | 2.77E-02 |
| positive regulation of cell growth | GO:0030307 | 99 | 4 | 0.26 | + | 15.41 | 1.53E-04 | 5.71E-03 |
| regulation of cell-matrix adhesion | GO:0001952 | 75 | 3 | 0.2 | + | 15.25 | 1.12E-03 | 2.83E-02 |
| cell fate specification | GO:0001708 | 75 | 3 | 0.2 | + | 15.25 | 1.12E-03 | 2.83E-02 |
| regulation of ossification | GO:0030278 | 150 | 6 | 0.39 | + | 15.25 | 3.27E-06 | 2.51E-04 |
| regulation of receptor signaling pathway via JAK-STAT | GO:0046425 | 100 | 4 | 0.26 | + | 15.25 | 1.59E-04 | 5.91E-03 |
| negative regulation of neurogenesis | GO:0050768 | 176 | 7 | 0.46 | + | 15.17 | 4.84E-07 | 5.02E-05 |
| epidermis development | GO:0008544 | 151 | 6 | 0.4 | + | 15.15 | 3.39E-06 | 2.59E-04 |

| | | | | | | | | |
|---|---|---|---|---|---|---|---|---|
| gonad development | GO:0008406 | 126 | 5 | 0.33 | + | 15.13 | 2.36E-05 | 1.24E-03 |
| regulation of receptor signaling pathway via STAT | GO:1904892 | 101 | 4 | 0.26 | + | 15.1 | 1.65E-04 | 6.11E-03 |
| male gonad development | GO:0008584 | 76 | 3 | 0.2 | + | 15.05 | 1.16E-03 | 2.92E-02 |
| development of primary male sexual characteristics | GO:0046546 | 77 | 3 | 0.2 | + | 14.86 | 1.21E-03 | 3.01E-02 |
| regulation of reactive oxygen species metabolic process | GO:2000377 | 103 | 4 | 0.27 | + | 14.81 | 1.77E-04 | 6.51E-03 |
| regulation of leukocyte proliferation | GO:0070663 | 155 | 6 | 0.41 | + | 14.76 | 3.92E-06 | 2.88E-04 |
| reactive oxygen species metabolic process | GO:0072593 | 78 | 3 | 0.2 | + | 14.67 | 1.25E-03 | 3.08E-02 |
| development of primary sexual characteristics | GO:0045137 | 130 | 5 | 0.34 | + | 14.67 | 2.73E-05 | 1.39E-03 |
| muscle cell development | GO:0055001 | 105 | 4 | 0.28 | + | 14.53 | 1.90E-04 | 6.86E-03 |
| epidermal cell differentiation | GO:0009913 | 79 | 3 | 0.21 | + | 14.48 | 1.30E-03 | 3.18E-02 |
| B cell differentiation | GO:0030183 | 79 | 3 | 0.21 | + | 14.48 | 1.30E-03 | 3.18E-02 |

| | | | | | | | | |
|---|---|---|---|---|---|---|---|---|
| aging | GO:0007568 | 79 | 3 | 0.21 | + | 14.48 | 1.30E-03 | 3.17E-02 |
| striated muscle tissue development | GO:0014706 | 211 | 8 | 0.55 | + | 14.46 | 9.82E-08 | 1.25E-05 |
| regulation of animal organ morphogenesis | GO:2000027 | 132 | 5 | 0.35 | + | 14.44 | 2.93E-05 | 1.45E-03 |
| female gamete generation | GO:0007292 | 80 | 3 | 0.21 | + | 14.3 | 1.34E-03 | 3.25E-02 |
| spinal cord development | GO:0021510 | 81 | 3 | 0.21 | + | 14.12 | 1.39E-03 | 3.33E-02 |
| regulation of neuron apoptotic process | GO:0043523 | 108 | 4 | 0.28 | + | 14.12 | 2.11E-04 | 7.50E-03 |
| mesoderm development | GO:0007498 | 82 | 3 | 0.22 | + | 13.95 | 1.44E-03 | 3.40E-02 |
| epithelial cell development | GO:0002064 | 137 | 5 | 0.36 | + | 13.92 | 3.48E-05 | 1.68E-03 |
| determination of left/right symmetry | GO:0007368 | 83 | 3 | 0.22 | + | 13.78 | 1.49E-03 | 3.49E-02 |
| myelination | GO:0042552 | 83 | 3 | 0.22 | + | 13.78 | 1.49E-03 | 3.49E-02 |
| negative regulation of nervous system development | GO:0051961 | 194 | 7 | 0.51 | + | 13.76 | 9.09E-07 | 8.39E-05 |

| | | | | | | | | |
|---|---|---|---|---|---|---|---|---|
| hindbrain development | GO:0030902 | 111 | 4 | 0.29 | + | 13.74 | 2.34E-04 | 8.14E-03 |
| cellular response to biotic stimulus | GO:0071216 | 140 | 5 | 0.37 | + | 13.62 | 3.85E-05 | 1.85E-03 |
| regulation of T cell proliferation | GO:0042129 | 112 | 4 | 0.29 | + | 13.62 | 2.42E-04 | 8.40E-03 |
| muscle tissue development | GO:0060537 | 224 | 8 | 0.59 | + | 13.62 | 1.53E-07 | 1.84E-05 |
| axon guidance | GO:0007411 | 169 | 6 | 0.44 | + | 13.54 | 6.33E-06 | 4.26E-04 |
| ensheathment of neurons | GO:0007272 | 85 | 3 | 0.22 | + | 13.46 | 1.59E-03 | 3.65E-02 |
| axon ensheathment | GO:0008366 | 85 | 3 | 0.22 | + | 13.46 | 1.59E-03 | 3.64E-02 |
| positive regulation of growth | GO:0045927 | 170 | 6 | 0.45 | + | 13.46 | 6.54E-06 | 4.36E-04 |
| transmembrane receptor protein serine/threonine kinase signaling pathway | GO:0007178 | 142 | 5 | 0.37 | + | 13.43 | 4.11E-05 | 1.96E-03 |
| muscle organ development | GO:0007517 | 171 | 6 | 0.45 | + | 13.38 | 6.76E-06 | 4.42E-04 |
| regulation of carbohydrate metabolic process | GO:0006109 | 114 | 4 | 0.3 | + | 13.38 | 2.58E-04 | 8.81E-03 |

| | | | | | | | | |
|---|---|---|---|---|---|---|---|---|
| mesenchymal cell differentiation | GO:0048762 | 114 | 4 | 0.3 | + | 13.38 | 2.58E-04 | 8.79E-03 |
| negative regulation of leukocyte cell-cell adhesion | GO:1903038 | 86 | 3 | 0.23 | + | 13.3 | 1.64E-03 | 3.72E-02 |
| regulation of muscle cell differentiation | GO:0051147 | 86 | 3 | 0.23 | + | 13.3 | 1.64E-03 | 3.72E-02 |
| regulation of DNA binding | GO:0051101 | 86 | 3 | 0.23 | + | 13.3 | 1.64E-03 | 3.71E-02 |
| neuron projection guidance | GO:0097485 | 172 | 6 | 0.45 | + | 13.3 | 6.98E-06 | 4.55E-04 |
| T cell differentiation | GO:0030217 | 115 | 4 | 0.3 | + | 13.26 | 2.67E-04 | 9.06E-03 |
| negative regulation of secretion by cell | GO:1903531 | 115 | 4 | 0.3 | + | 13.26 | 2.67E-04 | 9.04E-03 |
| positive regulation of transmembrane receptor protein serine/threonine kinase signaling pathway | GO:0090100 | 87 | 3 | 0.23 | + | 13.15 | 1.69E-03 | 3.83E-02 |
| regulation of striated muscle tissue development | GO:0016202 | 87 | 3 | 0.23 | + | 13.15 | 1.69E-03 | 3.82E-02 |
| cardiac muscle tissue development | GO:0048738 | 116 | 4 | 0.3 | + | 13.15 | 2.75E-04 | 9.22E-03 |
| determination of bilateral symmetry | GO:0009855 | 88 | 3 | 0.23 | + | 13 | 1.75E-03 | 3.93E-02 |

| | | | | | | | | |
|---|---|---|---|---|---|---|---|---|
| cellular response to lipopolysaccharide | GO:0071222 | 118 | 4 | 0.31 | + | 12.93 | 2.93E-04 | 9.77E-03 |
| peptidyl-tyrosine phosphorylation | GO:0018108 | 118 | 4 | 0.31 | + | 12.93 | 2.93E-04 | 9.75E-03 |
| adaptive immune response based on somatic recombination of immune receptors built from immunoglobulin superfamily domains | GO:0002460 | 89 | 3 | 0.23 | + | 12.85 | 1.80E-03 | 4.00E-02 |
| regulation of muscle tissue development | GO:1901861 | 89 | 3 | 0.23 | + | 12.85 | 1.80E-03 | 3.99E-02 |
| positive regulation of peptidyl-serine phosphorylation | GO:0033138 | 89 | 3 | 0.23 | + | 12.85 | 1.80E-03 | 3.99E-02 |
| specification of symmetry | GO:0009799 | 89 | 3 | 0.23 | + | 12.85 | 1.80E-03 | 3.98E-02 |
| cell surface receptor signaling pathway involved in cell-cell signaling | GO:1905114 | 178 | 6 | 0.47 | + | 12.85 | 8.44E-06 | 5.37E-04 |
| cellular response to growth factor stimulus | GO:0071363 | 298 | 10 | 0.78 | + | 12.8 | 6.51E-09 | 1.25E-06 |
| negative regulation of mitotic cell cycle phase transition | GO:1901991 | 90 | 3 | 0.24 | + | 12.71 | 1.86E-03 | 4.10E-02 |
| peptidyl-tyrosine modification | GO:0018212 | 120 | 4 | 0.31 | + | 12.71 | 3.12E-04 | 1.02E-02 |
| regulation of muscle organ development | GO:0048634 | 90 | 3 | 0.24 | + | 12.71 | 1.86E-03 | 4.09E-02 |

| | | | | | | | | |
|---|---|---|---|---|---|---|---|---|
| heart morphogenesis | GO:0003007 | 181 | 6 | 0.47 | + | 12.64 | 9.25E-06 | 5.74E-04 |
| anterior/posterior pattern specification | GO:0009952 | 182 | 6 | 0.48 | + | 12.57 | 9.54E-06 | 5.87E-04 |
| positive regulation of epithelial cell proliferation | GO:0050679 | 122 | 4 | 0.32 | + | 12.5 | 3.31E-04 | 1.07E-02 |
| cellular response to molecule of bacterial origin | GO:0071219 | 123 | 4 | 0.32 | + | 12.4 | 3.42E-04 | 1.11E-02 |
| response to growth factor | GO:0070848 | 308 | 10 | 0.81 | + | 12.38 | 8.82E-09 | 1.63E-06 |
| male sex differentiation | GO:0046661 | 93 | 3 | 0.24 | + | 12.3 | 2.04E-03 | 4.45E-02 |
| cardiac ventricle development | GO:0003231 | 93 | 3 | 0.24 | + | 12.3 | 2.04E-03 | 4.44E-02 |
| positive regulation of peptidyl-tyrosine phosphorylation | GO:0050731 | 124 | 4 | 0.33 | + | 12.3 | 3.52E-04 | 1.13E-02 |
| ossification | GO:0001503 | 156 | 5 | 0.41 | + | 12.22 | 6.33E-05 | 2.85E-03 |
| endocrine system development | GO:0035270 | 94 | 3 | 0.25 | + | 12.17 | 2.10E-03 | 4.51E-02 |
| axonogenesis | GO:0007409 | 251 | 8 | 0.66 | + | 12.15 | 3.54E-07 | 3.87E-05 |

| | | | | | | | | |
|---|---|---|---|---|---|---|---|---|
| homeostasis of number of cells | GO:0048872 | 159 | 5 | 0.42 | + | 11.99 | 6.91E-05 | 3.01E-03 |
| limb morphogenesis | GO:0035108 | 128 | 4 | 0.34 | + | 11.92 | 3.95E-04 | 1.25E-02 |
| appendage morphogenesis | GO:0035107 | 128 | 4 | 0.34 | + | 11.92 | 3.95E-04 | 1.24E-02 |
| stem cell differentiation | GO:0048863 | 129 | 4 | 0.34 | + | 11.82 | 4.07E-04 | 1.27E-02 |
| negative regulation of secretion | GO:0051048 | 129 | 4 | 0.34 | + | 11.82 | 4.07E-04 | 1.27E-02 |
| striated muscle cell development | GO:0055002 | 97 | 3 | 0.25 | + | 11.79 | 2.29E-03 | 4.85E-02 |
| sex differentiation | GO:0007548 | 163 | 5 | 0.43 | + | 11.7 | 7.75E-05 | 3.32E-03 |
| cell morphogenesis involved in neuron differentiation | GO:0048667 | 301 | 9 | 0.79 | + | 11.4 | 1.04E-07 | 1.31E-05 |
| regulation of cell growth | GO:0001558 | 235 | 7 | 0.62 | + | 11.36 | 3.13E-06 | 2.41E-04 |
| muscle cell differentiation | GO:0042692 | 168 | 5 | 0.44 | + | 11.35 | 8.90E-05 | 3.73E-03 |
| axon development | GO:0061564 | 270 | 8 | 0.71 | + | 11.3 | 6.05E-07 | 5.93E-05 |

| | | | | | | | | |
|---|---|---|---|---|---|---|---|---|
| positive regulation of cell population proliferation | GO:0008284 | 574 | 17 | 1.51 | + | 11.29 | 9.01E-14 | 1.60E-10 |
| response to peptide | GO:1901652 | 203 | 6 | 0.53 | + | 11.27 | 1.74E-05 | 9.58E-04 |
| negative regulation of neuron differentiation | GO:0045665 | 139 | 4 | 0.36 | + | 10.97 | 5.35E-04 | 1.59E-02 |
| negative regulation of cell migration | GO:0030336 | 176 | 5 | 0.46 | + | 10.83 | 1.10E-04 | 4.42E-03 |
| negative regulation of cell differentiation | GO:0045596 | 461 | 13 | 1.21 | + | 10.75 | 2.07E-10 | 9.48E-08 |
| regulation of cell-substrate adhesion | GO:0010810 | 142 | 4 | 0.37 | + | 10.74 | 5.79E-04 | 1.70E-02 |
| negative regulation of cell adhesion | GO:0007162 | 182 | 5 | 0.48 | + | 10.48 | 1.28E-04 | 5.02E-03 |
| epithelial cell differentiation | GO:0030855 | 328 | 9 | 0.86 | + | 10.46 | 2.11E-07 | 2.45E-05 |
| enzyme linked receptor protein signaling pathway | GO:0007167 | 439 | 12 | 1.15 | + | 10.42 | 1.59E-09 | 3.90E-07 |
| negative regulation of cell motility | GO:2000146 | 183 | 5 | 0.48 | + | 10.42 | 1.31E-04 | 5.13E-03 |
| heart development | GO:0007507 | 369 | 10 | 0.97 | + | 10.33 | 4.63E-08 | 6.71E-06 |

| | | | | | | | | |
|---|---|---|---|---|---|---|---|---|
| lymphocyte differentiation | GO:0030098 | 186 | 5 | 0.49 | + | 10.25 | 1.42E-04 | 5.38E-03 |
| limb development | GO:0060173 | 150 | 4 | 0.39 | + | 10.17 | 7.07E-04 | 2.00E-02 |
| appendage development | GO:0048736 | 150 | 4 | 0.39 | + | 10.17 | 7.07E-04 | 2.00E-02 |
| cytokine-mediated signaling pathway | GO:0019221 | 263 | 7 | 0.69 | + | 10.15 | 6.42E-06 | 4.30E-04 |
| gland development | GO:0048732 | 264 | 7 | 0.69 | + | 10.11 | 6.58E-06 | 4.37E-04 |
| muscle structure development | GO:0061061 | 305 | 8 | 0.8 | + | 10 | 1.47E-06 | 1.25E-04 |
| negative regulation of growth | GO:0045926 | 154 | 4 | 0.4 | + | 9.9 | 7.78E-04 | 2.18E-02 |
| positive regulation of cell migration | GO:0030335 | 348 | 9 | 0.91 | + | 9.86 | 3.42E-07 | 3.77E-05 |
| regulation of mitotic cell cycle phase transition | GO:1901990 | 195 | 5 | 0.51 | + | 9.78 | 1.76E-04 | 6.46E-03 |
| regulation of neuron death | GO:1901214 | 156 | 4 | 0.41 | + | 9.78 | 8.15E-04 | 2.23E-02 |
| regionalization | GO:0003002 | 274 | 7 | 0.72 | + | 9.74 | 8.33E-06 | 5.33E-04 |

| | | | | | | | | |
|---|---|---|---|---|---|---|---|---|
| cellular response to peptide | GO:1901653 | 158 | 4 | 0.41 | + | 9.65 | 8.54E-04 | 2.30E-02 |
| positive regulation of neurogenesis | GO:0050769 | 278 | 7 | 0.73 | + | 9.6 | 9.13E-06 | 5.69E-04 |
| mesenchyme development | GO:0060485 | 159 | 4 | 0.42 | + | 9.59 | 8.74E-04 | 2.35E-02 |
| negative regulation of cell population proliferation | GO:0008285 | 439 | 11 | 1.15 | + | 9.56 | 1.99E-08 | 3.25E-06 |
| regulation of leukocyte cell-cell adhesion | GO:1903037 | 200 | 5 | 0.52 | + | 9.53 | 1.97E-04 | 7.04E-03 |
| camera-type eye development | GO:0043010 | 240 | 6 | 0.63 | + | 9.53 | 4.35E-05 | 2.06E-03 |
| taxis | GO:0042330 | 361 | 9 | 0.95 | + | 9.51 | 4.62E-07 | 4.86E-05 |
| cell population proliferation | GO:0008283 | 361 | 9 | 0.95 | + | 9.51 | 4.62E-07 | 4.82E-05 |
| reproductive structure development | GO:0048608 | 282 | 7 | 0.74 | + | 9.47 | 1.00E-05 | 6.10E-04 |
| positive regulation of cell motility | GO:2000147 | 364 | 9 | 0.95 | + | 9.43 | 4.94E-07 | 5.05E-05 |
| reproductive system development | GO:0061458 | 284 | 7 | 0.74 | + | 9.4 | 1.05E-05 | 6.32E-04 |

| | | | | | | | | |
|---|---|---|---|---|---|---|---|---|
| regulation of protein stability | GO:0031647 | 204 | 5 | 0.53 | + | 9.35 | 2.15E-04 | 7.57E-03 |
| positive regulation of cellular component movement | GO:0051272 | 368 | 9 | 0.97 | + | 9.33 | 5.40E-07 | 5.48E-05 |
| neuron projection morphogenesis | GO:0048812 | 331 | 8 | 0.87 | + | 9.22 | 2.67E-06 | 2.16E-04 |
| developmental growth | GO:0048589 | 290 | 7 | 0.76 | + | 9.2 | 1.19E-05 | 7.04E-04 |
| plasma membrane bounded cell projection morphogenesis | GO:0120039 | 333 | 8 | 0.87 | + | 9.16 | 2.78E-06 | 2.23E-04 |
| response to lipopolysaccharide | GO:0032496 | 167 | 4 | 0.44 | + | 9.13 | 1.04E-03 | 2.68E-02 |
| growth | GO:0040007 | 293 | 7 | 0.77 | + | 9.11 | 1.27E-05 | 7.42E-04 |
| regulation of lymphocyte activation | GO:0051249 | 293 | 7 | 0.77 | + | 9.11 | 1.27E-05 | 7.39E-04 |
| cell projection morphogenesis | GO:0048858 | 335 | 8 | 0.88 | + | 9.11 | 2.91E-06 | 2.29E-04 |
| negative regulation of cellular component movement | GO:0051271 | 210 | 5 | 0.55 | + | 9.08 | 2.45E-04 | 8.51E-03 |
| regulation of small molecule metabolic process | GO:0062012 | 210 | 5 | 0.55 | + | 9.08 | 2.45E-04 | 8.48E-03 |

| | | | | | | | | |
|---|---|---|---|---|---|---|---|---|
| positive regulation of locomotion | GO:0040017 | 382 | 9 | 1 | + | 8.98 | 7.31E-07 | 7.07E-05 |
| inflammatory response | GO:0006954 | 255 | 6 | 0.67 | + | 8.97 | 6.04E-05 | 2.74E-03 |
| blood vessel morphogenesis | GO:0048514 | 256 | 6 | 0.67 | + | 8.94 | 6.17E-05 | 2.79E-03 |
| regulation of binding | GO:0051098 | 257 | 6 | 0.67 | + | 8.9 | 6.30E-05 | 2.84E-03 |
| regulation of peptidyl-tyrosine phosphorylation | GO:0050730 | 172 | 4 | 0.45 | + | 8.87 | 1.16E-03 | 2.92E-02 |
| regulation of leukocyte activation | GO:0002694 | 344 | 8 | 0.9 | + | 8.87 | 3.52E-06 | 2.67E-04 |
| regulation of growth | GO:0040008 | 389 | 9 | 1.02 | + | 8.82 | 8.48E-07 | 7.97E-05 |
| regulation of hemopoiesis | GO:1903706 | 260 | 6 | 0.68 | + | 8.8 | 6.70E-05 | 2.93E-03 |
| regulation of neurogenesis | GO:0050767 | 523 | 12 | 1.37 | + | 8.75 | 1.08E-08 | 1.91E-06 |
| regulation of vasculature development | GO:1901342 | 218 | 5 | 0.57 | + | 8.75 | 2.91E-04 | 9.71E-03 |
| regulation of cell population proliferation | GO:0042127 | 1047 | 24 | 2.75 | + | 8.74 | 4.39E-17 | 6.24E-13 |

| | | | | | | | | |
|---|---|---|---|---|---|---|---|---|
| negative regulation of developmental process | GO:0051093 | 656 | 15 | 1.72 | + | 8.72 | 1.17E-10 | 6.14E-08 |
| negative regulation of locomotion | GO:0040013 | 219 | 5 | 0.57 | + | 8.71 | 2.97E-04 | 9.84E-03 |
| cell part morphogenesis | GO:0032990 | 351 | 8 | 0.92 | + | 8.69 | 4.07E-06 | 2.93E-04 |
| regulation of cell cycle phase transition | GO:1901987 | 221 | 5 | 0.58 | + | 8.63 | 3.09E-04 | 1.01E-02 |
| cell morphogenesis involved in differentiation | GO:0000904 | 398 | 9 | 1.04 | + | 8.62 | 1.02E-06 | 9.06E-05 |
| response to molecule of bacterial origin | GO:0002237 | 179 | 4 | 0.47 | + | 8.52 | 1.34E-03 | 3.26E-02 |
| response to light stimulus | GO:0009416 | 179 | 4 | 0.47 | + | 8.52 | 1.34E-03 | 3.26E-02 |
| eye development | GO:0001654 | 269 | 6 | 0.71 | + | 8.51 | 8.05E-05 | 3.44E-03 |
| chemotaxis | GO:0006935 | 359 | 8 | 0.94 | + | 8.5 | 4.78E-06 | 3.33E-04 |
| visual system development | GO:0150063 | 270 | 6 | 0.71 | + | 8.47 | 8.22E-05 | 3.49E-03 |
| lymphocyte activation | GO:0046649 | 271 | 6 | 0.71 | + | 8.44 | 8.38E-05 | 3.55E-03 |

| | | | | | | | | |
|---|---|---|---|---|---|---|---|---|
| positive regulation of DNA-binding transcription factor activity | GO:0051091 | 181 | 4 | 0.47 | + | 8.43 | 1.40E-03 | 3.34E-02 |
| sensory system development | GO:0048880 | 272 | 6 | 0.71 | + | 8.41 | 8.55E-05 | 3.61E-03 |
| cellular response to cytokine stimulus | GO:0071345 | 501 | 11 | 1.31 | + | 8.37 | 7.40E-08 | 9.64E-06 |
| positive regulation of protein phosphorylation | GO:0001934 | 731 | 16 | 1.92 | + | 8.35 | 4.65E-11 | 3.67E-08 |
| positive regulation of lymphocyte activation | GO:0051251 | 183 | 4 | 0.48 | + | 8.34 | 1.45E-03 | 3.43E-02 |
| regulation of developmental growth | GO:0048638 | 229 | 5 | 0.6 | + | 8.33 | 3.62E-04 | 1.16E-02 |
| regulation of cell activation | GO:0050865 | 367 | 8 | 0.96 | + | 8.31 | 5.60E-06 | 3.83E-04 |
| T cell activation | GO:0042110 | 184 | 4 | 0.48 | + | 8.29 | 1.48E-03 | 3.48E-02 |
| regulation of transmembrane receptor protein serine/threonine kinase signaling pathway | GO:0090092 | 185 | 4 | 0.49 | + | 8.25 | 1.51E-03 | 3.48E-02 |
| regulation of cell migration | GO:0030334 | 604 | 13 | 1.58 | + | 8.21 | 5.05E-09 | 1.04E-06 |
| response to cytokine | GO:0034097 | 558 | 12 | 1.46 | + | 8.2 | 2.17E-08 | 3.50E-06 |

| | | | | | | | | |
|---|---|---|---|---|---|---|---|---|
| regulation of apoptotic signaling pathway | GO:2001233 | 279 | 6 | 0.73 | + | 8.2 | 9.80E-05 | 4.05E-03 |
| sensory organ development | GO:0007423 | 419 | 9 | 1.1 | + | 8.19 | 1.55E-06 | 1.31E-04 |
| embryonic organ morphogenesis | GO:0048562 | 233 | 5 | 0.61 | + | 8.18 | 3.92E-04 | 1.24E-02 |
| tube development | GO:0035295 | 606 | 13 | 1.59 | + | 8.18 | 5.25E-09 | 1.05E-06 |
| regulation of cell development | GO:0060284 | 610 | 13 | 1.6 | + | 8.13 | 5.67E-09 | 1.10E-06 |
| regulation of leukocyte differentiation | GO:1902105 | 188 | 4 | 0.49 | + | 8.11 | 1.60E-03 | 3.66E-02 |
| positive regulation of cell development | GO:0010720 | 331 | 7 | 0.87 | + | 8.06 | 2.74E-05 | 1.39E-03 |
| regulation of cell adhesion | GO:0030155 | 473 | 10 | 1.24 | + | 8.06 | 4.37E-07 | 4.63E-05 |
| regulation of DNA-binding transcription factor activity | GO:0051090 | 284 | 6 | 0.74 | + | 8.06 | 1.08E-04 | 4.35E-03 |
| positive regulation of nervous system development | GO:0051962 | 334 | 7 | 0.88 | + | 7.99 | 2.90E-05 | 1.44E-03 |
| cell activation | GO:0001775 | 383 | 8 | 1 | + | 7.97 | 7.61E-06 | 4.89E-04 |

| | | | | | | | | |
|---|---|---|---|---|---|---|---|---|
| cellular response to lipid | GO:0071396 | 288 | 6 | 0.76 | + | 7.94 | 1.16E-04 | 4.61E-03 |
| pattern specification process | GO:0007389 | 340 | 7 | 0.89 | + | 7.85 | 3.24E-05 | 1.58E-03 |
| response to lipid | GO:0033993 | 389 | 8 | 1.02 | + | 7.84 | 8.50E-06 | 5.36E-04 |
| transmembrane receptor protein tyrosine kinase signaling pathway | GO:0007169 | 292 | 6 | 0.77 | + | 7.84 | 1.25E-04 | 4.91E-03 |
| circulatory system development | GO:0072359 | 588 | 12 | 1.54 | + | 7.78 | 3.81E-08 | 5.58E-06 |
| innate immune response | GO:0045087 | 245 | 5 | 0.64 | + | 7.78 | 4.90E-04 | 1.48E-02 |
| negative regulation of response to external stimulus | GO:0032102 | 245 | 5 | 0.64 | + | 7.78 | 4.90E-04 | 1.48E-02 |
| epithelium development | GO:0060429 | 688 | 14 | 1.8 | + | 7.76 | 2.37E-09 | 5.34E-07 |
| regulation of cell motility | GO:2000145 | 641 | 13 | 1.68 | + | 7.73 | 1.01E-08 | 1.82E-06 |
| positive regulation of phosphorylation | GO:0042327 | 789 | 16 | 2.07 | + | 7.73 | 1.40E-10 | 6.87E-08 |
| regulation of angiogenesis | GO:0045765 | 199 | 4 | 0.52 | + | 7.67 | 1.96E-03 | 4.28E-02 |

| | | | | | | | | |
|---|---|---|---|---|---|---|---|---|
| tube morphogenesis | GO:0035239 | 451 | 9 | 1.18 | + | 7.61 | 2.79E-06 | 2.23E-04 |
| response to endogenous stimulus | GO:0009719 | 819 | 16 | 2.15 | + | 7.45 | 2.40E-10 | 1.03E-07 |
| positive regulation of leukocyte activation | GO:0002696 | 206 | 4 | 0.54 | + | 7.4 | 2.22E-03 | 4.75E-02 |
| regulation of nervous system development | GO:0051960 | 620 | 12 | 1.63 | + | 7.38 | 6.71E-08 | 8.83E-06 |
| regulation of T cell activation | GO:0050863 | 207 | 4 | 0.54 | + | 7.37 | 2.26E-03 | 4.83E-02 |
| leukocyte differentiation | GO:0002521 | 260 | 5 | 0.68 | + | 7.33 | 6.38E-04 | 1.84E-02 |
| positive regulation of phosphorus metabolic process | GO:0010562 | 833 | 16 | 2.18 | + | 7.32 | 3.06E-10 | 1.24E-07 |
| positive regulation of phosphate metabolic process | GO:0045937 | 833 | 16 | 2.18 | + | 7.32 | 3.06E-10 | 1.21E-07 |
| positive regulation of MAPK cascade | GO:0043410 | 366 | 7 | 0.96 | + | 7.29 | 5.12E-05 | 2.38E-03 |
| regulation of neuron differentiation | GO:0045664 | 422 | 8 | 1.11 | + | 7.23 | 1.51E-05 | 8.50E-04 |
| positive regulation of cell differentiation | GO:0045597 | 635 | 12 | 1.67 | + | 7.21 | 8.66E-08 | 1.12E-05 |

| | | | | | | | | |
|---|---|---|---|---|---|---|---|---|
| regulation of cellular component movement | GO:0051270 | 693 | 13 | 1.82 | + | 7.15 | 2.50E-08 | 3.98E-06 |
| negative regulation of multicellular organismal process | GO:0051241 | 800 | 15 | 2.1 | + | 7.15 | 1.70E-09 | 4.09E-07 |
| regulation of locomotion | GO:0040012 | 694 | 13 | 1.82 | + | 7.14 | 2.54E-08 | 4.01E-06 |
| blood vessel development | GO:0001568 | 324 | 6 | 0.85 | + | 7.06 | 2.17E-04 | 7.63E-03 |
| leukocyte activation | GO:0045321 | 325 | 6 | 0.85 | + | 7.04 | 2.21E-04 | 7.73E-03 |
| regulation of cell-cell adhesion | GO:0022407 | 271 | 5 | 0.71 | + | 7.04 | 7.66E-04 | 2.15E-02 |
| cell surface receptor signaling pathway | GO:0007166 | 1367 | 25 | 3.58 | + | 6.97 | 1.35E-15 | 6.39E-12 |
| positive regulation of protein modification process | GO:0031401 | 894 | 16 | 2.34 | + | 6.82 | 8.38E-10 | 2.38E-07 |
| regulation of MAPK cascade | GO:0043408 | 503 | 9 | 1.32 | + | 6.82 | 6.65E-06 | 4.37E-04 |
| neuron development | GO:0048666 | 561 | 10 | 1.47 | + | 6.8 | 1.99E-06 | 1.65E-04 |
| embryonic organ development | GO:0048568 | 337 | 6 | 0.88 | + | 6.79 | 2.68E-04 | 9.05E-03 |

| | | | | | | | | |
|---|---|---|---|---|---|---|---|---|
| cellular response to organic substance | GO:0071310 | 1405 | 25 | 3.68 | + | 6.79 | 2.51E-15 | 8.93E-12 |
| embryonic morphogenesis | GO:0048598 | 450 | 8 | 1.18 | + | 6.78 | 2.38E-05 | 1.24E-03 |
| immune effector process | GO:0002252 | 282 | 5 | 0.74 | + | 6.76 | 9.13E-04 | 2.43E-02 |
| defense response | GO:0006952 | 680 | 12 | 1.78 | + | 6.73 | 1.79E-07 | 2.10E-05 |
| cellular response to endogenous stimulus | GO:0071495 | 743 | 13 | 1.95 | + | 6.67 | 5.56E-08 | 7.52E-06 |
| neuron projection development | GO:0031175 | 459 | 8 | 1.2 | + | 6.65 | 2.74E-05 | 1.39E-03 |
| negative regulation of transport | GO:0051051 | 287 | 5 | 0.75 | + | 6.64 | 9.86E-04 | 2.59E-02 |
| vasculature development | GO:0001944 | 345 | 6 | 0.9 | + | 6.63 | 3.03E-04 | 9.98E-03 |
| generation of neurons | GO:0048699 | 1036 | 18 | 2.72 | + | 6.63 | 8.97E-11 | 5.10E-08 |
| central nervous system development | GO:0007417 | 578 | 10 | 1.52 | + | 6.6 | 2.58E-06 | 2.11E-04 |
| negative regulation of protein phosphorylation | GO:0001933 | 289 | 5 | 0.76 | + | 6.6 | 1.02E-03 | 2.64E-02 |

| | | | | | | | | |
|---|---|---|---|---|---|---|---|---|
| positive regulation of developmental process | GO:0051094 | 927 | 16 | 2.43 | + | 6.58 | 1.40E-09 | 3.56E-07 |
| negative regulation of immune system process | GO:0002683 | 290 | 5 | 0.76 | + | 6.57 | 1.03E-03 | 2.67E-02 |
| cell morphogenesis | GO:0000902 | 523 | 9 | 1.37 | + | 6.56 | 9.04E-06 | 5.66E-04 |
| neuron differentiation | GO:0030182 | 702 | 12 | 1.84 | + | 6.52 | 2.51E-07 | 2.87E-05 |
| cardiovascular system development | GO:0072358 | 354 | 6 | 0.93 | + | 6.46 | 3.46E-04 | 1.12E-02 |
| negative regulation of intracellular signal transduction | GO:1902532 | 358 | 6 | 0.94 | + | 6.39 | 3.67E-04 | 1.17E-02 |
| negative regulation of apoptotic process | GO:0043066 | 546 | 9 | 1.43 | + | 6.29 | 1.27E-05 | 7.41E-04 |
| tissue development | GO:0009888 | 1162 | 19 | 3.05 | + | 6.24 | 6.44E-11 | 4.36E-08 |
| cell junction organization | GO:0034330 | 306 | 5 | 0.8 | + | 6.23 | 1.30E-03 | 3.18E-02 |
| negative regulation of cell death | GO:0060548 | 612 | 10 | 1.6 | + | 6.23 | 4.25E-06 | 3.05E-04 |
| brain development | GO:0007420 | 429 | 7 | 1.12 | + | 6.22 | 1.36E-04 | 5.25E-03 |

| | | | | | | | | |
|---|---|---|---|---|---|---|---|---|
| cell-cell signaling | GO:0007267 | 553 | 9 | 1.45 | + | 6.21 | 1.40E-05 | 7.99E-04 |
| regulation of neuron projection development | GO:0010975 | 308 | 5 | 0.81 | + | 6.19 | 1.34E-03 | 3.25E-02 |
| regulation of cell differentiation | GO:0045595 | 1173 | 19 | 3.08 | + | 6.18 | 7.55E-11 | 4.88E-08 |
| response to oxygen-containing compound | GO:1901700 | 805 | 13 | 2.11 | + | 6.16 | 1.39E-07 | 1.68E-05 |
| neurogenesis | GO:0022008 | 1118 | 18 | 2.93 | + | 6.14 | 3.02E-10 | 1.26E-07 |
| in utero embryonic development | GO:0001701 | 311 | 5 | 0.82 | + | 6.13 | 1.40E-03 | 3.34E-02 |
| negative regulation of programmed cell death | GO:0043069 | 560 | 9 | 1.47 | + | 6.13 | 1.54E-05 | 8.64E-04 |
| regulation of protein phosphorylation | GO:0001932 | 1070 | 17 | 2.81 | + | 6.06 | 1.31E-09 | 3.38E-07 |
| protein phosphorylation | GO:0006468 | 693 | 11 | 1.82 | + | 6.05 | 1.74E-06 | 1.45E-04 |
| negative regulation of phosphorylation | GO:0042326 | 317 | 5 | 0.83 | + | 6.01 | 1.52E-03 | 3.50E-02 |
| cell development | GO:0048468 | 1146 | 18 | 3.01 | + | 5.99 | 4.47E-10 | 1.55E-07 |

| | | | | | | | | |
|---|---|---|---|---|---|---|---|---|
| positive regulation of cellular protein metabolic process | GO:0032270 | 1149 | 18 | 3.01 | + | 5.97 | 4.66E-10 | 1.58E-07 |
| response to organonitrogen compound | GO:0010243 | 511 | 8 | 1.34 | + | 5.97 | 5.77E-05 | 2.66E-03 |
| cellular response to chemical stimulus | GO:0070887 | 1793 | 28 | 4.7 | + | 5.96 | 6.39E-16 | 4.54E-12 |
| regulation of cytokine production | GO:0001817 | 449 | 7 | 1.18 | + | 5.95 | 1.79E-04 | 6.55E-03 |
| positive regulation of apoptotic process | GO:0043065 | 386 | 6 | 1.01 | + | 5.93 | 5.44E-04 | 1.61E-02 |
| positive regulation of programmed cell death | GO:0043068 | 389 | 6 | 1.02 | + | 5.88 | 5.66E-04 | 1.67E-02 |
| positive regulation of signal transduction | GO:0009967 | 1112 | 17 | 2.92 | + | 5.83 | 2.32E-09 | 5.32E-07 |
| animal organ morphogenesis | GO:0009887 | 720 | 11 | 1.89 | + | 5.83 | 2.50E-06 | 2.05E-04 |
| regulation of response to external stimulus | GO:0032101 | 590 | 9 | 1.55 | + | 5.82 | 2.32E-05 | 1.22E-03 |
| regulation of cellular response to stress | GO:0080135 | 460 | 7 | 1.21 | + | 5.8 | 2.07E-04 | 7.36E-03 |
| chordate embryonic development | GO:0043009 | 526 | 8 | 1.38 | + | 5.8 | 7.04E-05 | 3.05E-03 |

| | | | | | | | | |
|---|---|---|---|---|---|---|---|---|
| head development | GO:0060322 | 463 | 7 | 1.21 | + | 5.77 | 2.15E-04 | 7.58E-03 |
| behavior | GO:0007610 | 398 | 6 | 1.04 | + | 5.75 | 6.37E-04 | 1.84E-02 |
| negative regulation of transcription by RNA polymerase II | GO:0000122 | 664 | 10 | 1.74 | + | 5.74 | 8.61E-06 | 5.41E-04 |
| cellular component morphogenesis | GO:0032989 | 598 | 9 | 1.57 | + | 5.74 | 2.57E-05 | 1.32E-03 |
| embryo development ending in birth or egg hatching | GO:0009792 | 532 | 8 | 1.4 | + | 5.73 | 7.61E-05 | 3.28E-03 |
| response to organic substance | GO:0010033 | 1732 | 26 | 4.54 | + | 5.72 | 2.92E-14 | 8.30E-11 |
| cellular response to oxygen-containing compound | GO:1901701 | 604 | 9 | 1.58 | + | 5.68 | 2.78E-05 | 1.40E-03 |
| immune response | GO:0006955 | 672 | 10 | 1.76 | + | 5.67 | 9.54E-06 | 5.89E-04 |
| morphogenesis of an epithelium | GO:0002009 | 336 | 5 | 0.88 | + | 5.67 | 1.95E-03 | 4.28E-02 |
| positive regulation of transferase activity | GO:0051347 | 475 | 7 | 1.25 | + | 5.62 | 2.51E-04 | 8.65E-03 |
| positive regulation of cell death | GO:0010942 | 408 | 6 | 1.07 | + | 5.61 | 7.23E-04 | 2.04E-02 |

| | | | | | | | | |
|---|---|---|---|---|---|---|---|---|
| positive regulation of protein metabolic process | GO:0051247 | 1235 | 18 | 3.24 | + | 5.56 | 1.45E-09 | 3.62E-07 |
| response to hormone | GO:0009725 | 415 | 6 | 1.09 | + | 5.51 | 7.89E-04 | 2.17E-02 |
| negative regulation of phosphorus metabolic process | GO:0010563 | 415 | 6 | 1.09 | + | 5.51 | 7.89E-04 | 2.17E-02 |
| negative regulation of phosphate metabolic process | GO:0045936 | 415 | 6 | 1.09 | + | 5.51 | 7.89E-04 | 2.16E-02 |
| regulation of phosphorylation | GO:0042325 | 1180 | 17 | 3.09 | + | 5.49 | 5.59E-09 | 1.10E-06 |
| positive regulation of gene expression | GO:0010628 | 1528 | 22 | 4.01 | + | 5.49 | 1.38E-11 | 1.23E-08 |
| positive regulation of transcription by RNA polymerase II | GO:0045944 | 975 | 14 | 2.56 | + | 5.48 | 1.74E-07 | 2.05E-05 |
| positive regulation of kinase activity | GO:0033674 | 420 | 6 | 1.1 | + | 5.45 | 8.39E-04 | 2.28E-02 |
| regulation of anatomical structure morphogenesis | GO:0022603 | 703 | 10 | 1.84 | + | 5.42 | 1.40E-05 | 7.97E-04 |
| locomotion | GO:0040011 | 845 | 12 | 2.22 | + | 5.42 | 1.72E-06 | 1.45E-04 |
| positive regulation of transcription, DNA-templated | GO:0045893 | 1218 | 17 | 3.19 | + | 5.32 | 8.91E-09 | 1.62E-06 |

| | | | | | | | | |
|---|---|---|---|---|---|---|---|---|
| positive regulation of multicellular organismal process | GO:0051240 | 1148 | 16 | 3.01 | + | 5.31 | 2.78E-08 | 4.30E-06 |
| positive regulation of response to stimulus | GO:0048584 | 1507 | 21 | 3.95 | + | 5.31 | 8.72E-11 | 5.16E-08 |
| hemopoiesis | GO:0030097 | 431 | 6 | 1.13 | + | 5.31 | 9.57E-04 | 2.53E-02 |
| response to nitrogen compound | GO:1901698 | 577 | 8 | 1.51 | + | 5.29 | 1.32E-04 | 5.16E-03 |
| regulation of apoptotic process | GO:0042981 | 940 | 13 | 2.46 | + | 5.27 | 7.90E-07 | 7.53E-05 |
| regulation of multicellular organismal development | GO:2000026 | 1377 | 19 | 3.61 | + | 5.26 | 1.09E-09 | 2.92E-07 |
| positive regulation of macromolecule biosynthetic process | GO:0010557 | 1452 | 20 | 3.81 | + | 5.25 | 3.51E-10 | 1.35E-07 |
| defense response to other organism | GO:0098542 | 437 | 6 | 1.15 | + | 5.24 | 1.03E-03 | 2.66E-02 |
| phosphorylation | GO:0016310 | 952 | 13 | 2.5 | + | 5.21 | 9.09E-07 | 8.33E-05 |
| regulation of cell death | GO:0010941 | 1026 | 14 | 2.69 | + | 5.2 | 3.20E-07 | 3.55E-05 |
| regulation of programmed cell death | GO:0043067 | 956 | 13 | 2.51 | + | 5.19 | 9.53E-07 | 8.56E-05 |

| | | | | | | | | |
|---|---|---|---|---|---|---|---|---|
| positive regulation of cell communication | GO:0010647 | 1256 | 17 | 3.29 | + | 5.16 | 1.40E-08 | 2.39E-06 |
| positive regulation of intracellular signal transduction | GO:1902533 | 666 | 9 | 1.75 | + | 5.15 | 5.87E-05 | 2.70E-03 |
| positive regulation of signaling | GO:0023056 | 1261 | 17 | 3.31 | + | 5.14 | 1.48E-08 | 2.50E-06 |
| multicellular organismal reproductive process | GO:0048609 | 450 | 6 | 1.18 | + | 5.08 | 1.19E-03 | 2.98E-02 |
| positive regulation of cellular biosynthetic process | GO:0031328 | 1502 | 20 | 3.94 | + | 5.08 | 6.32E-10 | 1.95E-07 |
| positive regulation of RNA biosynthetic process | GO:1902680 | 1277 | 17 | 3.35 | + | 5.08 | 1.78E-08 | 2.97E-06 |
| positive regulation of nucleic acid-templated transcription | GO:1903508 | 1277 | 17 | 3.35 | + | 5.08 | 1.78E-08 | 2.94E-06 |
| response to abiotic stimulus | GO:0009628 | 601 | 8 | 1.58 | + | 5.08 | 1.74E-04 | 6.44E-03 |
| multi-organism reproductive process | GO:0044703 | 526 | 7 | 1.38 | + | 5.07 | 4.60E-04 | 1.39E-02 |
| positive regulation of biosynthetic process | GO:0009891 | 1520 | 20 | 3.99 | + | 5.02 | 7.76E-10 | 2.30E-07 |
| negative regulation of cell communication | GO:0010648 | 914 | 12 | 2.4 | + | 5.01 | 3.84E-06 | 2.84E-04 |

| | | | | | | | | |
|---|---|---|---|---|---|---|---|---|
| negative regulation of signaling | GO:0023057 | 917 | 12 | 2.4 | + | 4.99 | 3.97E-06 | 2.90E-04 |
| multicellular organism reproduction | GO:0032504 | 459 | 6 | 1.2 | + | 4.98 | 1.32E-03 | 3.21E-02 |
| multi-organism process | GO:0051704 | 536 | 7 | 1.41 | + | 4.98 | 5.13E-04 | 1.54E-02 |
| anatomical structure formation involved in morphogenesis | GO:0048646 | 619 | 8 | 1.62 | + | 4.93 | 2.13E-04 | 7.53E-03 |
| sexual reproduction | GO:0019953 | 466 | 6 | 1.22 | + | 4.91 | 1.42E-03 | 3.37E-02 |
| negative regulation of response to stimulus | GO:0048585 | 1088 | 14 | 2.85 | + | 4.91 | 6.43E-07 | 6.25E-05 |
| regulation of phosphate metabolic process | GO:0019220 | 1325 | 17 | 3.47 | + | 4.89 | 3.04E-08 | 4.65E-06 |
| regulation of phosphorus metabolic process | GO:0051174 | 1325 | 17 | 3.47 | + | 4.89 | 3.04E-08 | 4.60E-06 |
| regulation of response to stress | GO:0080134 | 858 | 11 | 2.25 | + | 4.89 | 1.29E-05 | 7.45E-04 |
| regulation of secretion by cell | GO:1903530 | 471 | 6 | 1.24 | + | 4.86 | 1.50E-03 | 3.50E-02 |
| hematopoietic or lymphoid organ development | GO:0048534 | 472 | 6 | 1.24 | + | 4.85 | 1.52E-03 | 3.49E-02 |

| | | | | | | | | |
|---|---|---|---|---|---|---|---|---|
| nervous system development | GO:0007399 | 1497 | 19 | 3.93 | + | 4.84 | 4.28E-09 | 9.07E-07 |
| positive regulation of RNA metabolic process | GO:0051254 | 1341 | 17 | 3.52 | + | 4.83 | 3.62E-08 | 5.36E-06 |
| regulation of transferase activity | GO:0051338 | 715 | 9 | 1.87 | + | 4.8 | 1.00E-04 | 4.10E-03 |
| regulation of intracellular signal transduction | GO:1902531 | 1114 | 14 | 2.92 | + | 4.79 | 8.49E-07 | 7.94E-05 |
| developmental process involved in reproduction | GO:0003006 | 559 | 7 | 1.47 | + | 4.78 | 6.56E-04 | 1.88E-02 |
| regulation of developmental process | GO:0050793 | 1770 | 22 | 4.64 | + | 4.74 | 2.31E-10 | 1.03E-07 |
| response to drug | GO:0042493 | 483 | 6 | 1.27 | + | 4.74 | 1.70E-03 | 3.83E-02 |
| regulation of immune system process | GO:0002682 | 891 | 11 | 2.34 | + | 4.71 | 1.83E-05 | 9.98E-04 |
| regulation of protein modification process | GO:0031399 | 1378 | 17 | 3.61 | + | 4.7 | 5.36E-08 | 7.39E-06 |
| anatomical structure morphogenesis | GO:0009653 | 1564 | 19 | 4.1 | + | 4.63 | 8.69E-09 | 1.62E-06 |
| immune system development | GO:0002520 | 501 | 6 | 1.31 | + | 4.57 | 2.04E-03 | 4.44E-02 |

| | | | | | | | | |
|---|---|---|---|---|---|---|---|---|
| regulation of secretion | GO:0051046 | 501 | 6 | 1.31 | + | 4.57 | 2.04E-03 | 4.43E-02 |
| negative regulation of signal transduction | GO:0009968 | 842 | 10 | 2.21 | + | 4.53 | 6.38E-05 | 2.86E-03 |
| positive regulation of macromolecule metabolic process | GO:0010604 | 2547 | 30 | 6.68 | + | 4.49 | 6.18E-14 | 1.25E-10 |
| regulation of signal transduction | GO:0009966 | 2055 | 24 | 5.39 | + | 4.45 | 8.67E-11 | 5.35E-08 |
| positive regulation of nucleobase-containing compound metabolic process | GO:0045935 | 1456 | 17 | 3.82 | + | 4.45 | 1.18E-07 | 1.47E-05 |
| animal organ development | GO:0048513 | 2064 | 24 | 5.41 | + | 4.43 | 9.48E-11 | 5.18E-08 |
| positive regulation of nitrogen compound metabolic process | GO:0051173 | 2415 | 28 | 6.33 | + | 4.42 | 1.06E-12 | 1.26E-09 |
| immune system process | GO:0002376 | 1294 | 15 | 3.39 | + | 4.42 | 8.67E-07 | 8.05E-05 |
| regulation of multicellular organismal process | GO:0051239 | 2085 | 24 | 5.47 | + | 4.39 | 1.17E-10 | 5.93E-08 |
| embryo development | GO:0009790 | 786 | 9 | 2.06 | + | 4.37 | 2.03E-04 | 7.24E-03 |
| positive regulation of cellular metabolic process | GO:0031325 | 2537 | 29 | 6.65 | + | 4.36 | 4.58E-13 | 5.92E-10 |

| | | | | | | | | |
|---|---|---|---|---|---|---|---|---|
| cell migration | GO:0016477 | 614 | 7 | 1.61 | + | 4.35 | 1.13E-03 | 2.84E-02 |
| intracellular signal transduction | GO:0035556 | 1142 | 13 | 2.99 | + | 4.34 | 6.60E-06 | 4.36E-04 |
| positive regulation of molecular function | GO:0044093 | 1238 | 14 | 3.25 | + | 4.31 | 2.91E-06 | 2.28E-04 |
| positive regulation of metabolic process | GO:0009893 | 2754 | 31 | 7.22 | + | 4.29 | 5.99E-14 | 1.42E-10 |
| response to external stimulus | GO:0009605 | 1335 | 15 | 3.5 | + | 4.28 | 1.28E-06 | 1.10E-04 |
| negative regulation of transcription, DNA-templated | GO:0045892 | 897 | 10 | 2.35 | + | 4.25 | 1.07E-04 | 4.34E-03 |
| negative regulation of RNA metabolic process | GO:0051253 | 998 | 11 | 2.62 | + | 4.2 | 5.10E-05 | 2.38E-03 |
| regulation of kinase activity | GO:0043549 | 642 | 7 | 1.68 | + | 4.16 | 1.45E-03 | 3.43E-02 |
| regulation of cell communication | GO:0010646 | 2401 | 26 | 6.3 | + | 4.13 | 4.99E-11 | 3.73E-08 |
| regulation of signaling | GO:0023051 | 2416 | 26 | 6.34 | + | 4.1 | 5.74E-11 | 4.07E-08 |
| response to stress | GO:0006950 | 2046 | 22 | 5.37 | + | 4.1 | 3.48E-09 | 7.73E-07 |

| | | | | | | | | |
|---|---|---|---|---|---|---|---|---|
| negative regulation of RNA biosynthetic process | GO:1902679 | 930 | 10 | 2.44 | + | 4.1 | 1.44E-04 | 5.41E-03 |
| negative regulation of nucleic acid-templated transcription | GO:1903507 | 930 | 10 | 2.44 | + | 4.1 | 1.44E-04 | 5.40E-03 |
| movement of cell or subcellular component | GO:0006928 | 1035 | 11 | 2.71 | + | 4.05 | 7.06E-05 | 3.05E-03 |
| regulation of response to stimulus | GO:0048583 | 2733 | 29 | 7.17 | + | 4.05 | 2.99E-12 | 3.27E-09 |
| plasma membrane bounded cell projection organization | GO:0120036 | 756 | 8 | 1.98 | + | 4.04 | 7.91E-04 | 2.16E-02 |
| regulation of transcription by RNA polymerase II | GO:0006357 | 1634 | 17 | 4.28 | + | 3.97 | 5.98E-07 | 5.98E-05 |
| cell projection organization | GO:0030030 | 770 | 8 | 2.02 | + | 3.96 | 8.89E-04 | 2.38E-02 |
| cell differentiation | GO:0030154 | 2325 | 24 | 6.1 | + | 3.94 | 1.08E-09 | 2.94E-07 |
| negative regulation of nucleobase-containing compound metabolic process | GO:0045934 | 1070 | 11 | 2.81 | + | 3.92 | 9.48E-05 | 3.96E-03 |
| positive regulation of catalytic activity | GO:0043085 | 997 | 10 | 2.61 | + | 3.82 | 2.51E-04 | 8.63E-03 |
| localization of cell | GO:0051674 | 699 | 7 | 1.83 | + | 3.82 | 2.34E-03 | 4.95E-02 |

| | | | | | | | | |
|---|---|---|---|---|---|---|---|---|
| cell motility | GO:0048870 | 699 | 7 | 1.83 | + | 3.82 | 2.34E-03 | 4.94E-02 |
| cellular developmental process | GO:0048869 | 2404 | 24 | 6.3 | + | 3.81 | 2.10E-09 | 4.98E-07 |
| negative regulation of molecular function | GO:0044092 | 816 | 8 | 2.14 | + | 3.74 | 1.29E-03 | 3.16E-02 |
| negative regulation of cellular macromolecule biosynthetic process | GO:2000113 | 1027 | 10 | 2.69 | + | 3.71 | 3.17E-04 | 1.03E-02 |
| regulation of cellular protein metabolic process | GO:0032268 | 1952 | 19 | 5.12 | + | 3.71 | 2.88E-07 | 3.22E-05 |
| reproductive process | GO:0022414 | 827 | 8 | 2.17 | + | 3.69 | 1.40E-03 | 3.34E-02 |
| reproduction | GO:0000003 | 829 | 8 | 2.17 | + | 3.68 | 1.42E-03 | 3.36E-02 |
| regulation of molecular function | GO:0065009 | 2180 | 21 | 5.72 | + | 3.67 | 6.22E-08 | 8.25E-06 |
| multicellular organism development | GO:0007275 | 3147 | 30 | 8.25 | + | 3.64 | 1.52E-11 | 1.27E-08 |
| regulation of localization | GO:0032879 | 1891 | 18 | 4.96 | + | 3.63 | 9.15E-07 | 8.34E-05 |
| system development | GO:0048731 | 2850 | 27 | 7.47 | + | 3.61 | 3.62E-10 | 1.35E-07 |

| | | | | | | | | |
|---|---|---|---|---|---|---|---|---|
| phosphate-containing compound metabolic process | GO:0006796 | 1587 | 15 | 4.16 | + | 3.6 | 1.04E-05 | 6.34E-04 |
| homeostatic process | GO:0042592 | 1174 | 11 | 3.08 | + | 3.57 | 2.13E-04 | 7.52E-03 |
| phosphorus metabolic process | GO:0006793 | 1608 | 15 | 4.22 | + | 3.56 | 1.22E-05 | 7.17E-04 |
| negative regulation of macromolecule biosynthetic process | GO:0010558 | 1075 | 10 | 2.82 | + | 3.55 | 4.54E-04 | 1.38E-02 |
| anatomical structure development | GO:0048856 | 3458 | 32 | 9.07 | + | 3.53 | 4.00E-12 | 4.06E-09 |
| regulation of protein metabolic process | GO:0051246 | 2080 | 19 | 5.45 | + | 3.48 | 7.62E-07 | 7.31E-05 |
| negative regulation of cellular metabolic process | GO:0031324 | 1878 | 17 | 4.92 | + | 3.45 | 4.01E-06 | 2.91E-04 |
| negative regulation of cellular biosynthetic process | GO:0031327 | 1111 | 10 | 2.91 | + | 3.43 | 5.86E-04 | 1.72E-02 |
| negative regulation of gene expression | GO:0010629 | 1227 | 11 | 3.22 | + | 3.42 | 3.10E-04 | 1.02E-02 |
| positive regulation of cellular process | GO:0048522 | 3912 | 35 | 10.26 | + | 3.41 | 3.88E-13 | 5.51E-10 |
| negative regulation of biosynthetic process | GO:0009890 | 1126 | 10 | 2.95 | + | 3.39 | 6.50E-04 | 1.87E-02 |

| | | | | | | | | |
|---|---|---|---|---|---|---|---|---|
| developmental process | GO:0032502 | 3720 | 33 | 9.76 | + | 3.38 | 4.46E-12 | 4.22E-09 |
| negative regulation of nitrogen compound metabolic process | GO:0051172 | 1747 | 15 | 4.58 | + | 3.27 | 3.23E-05 | 1.58E-03 |
| regulation of transport | GO:0051049 | 1168 | 10 | 3.06 | + | 3.26 | 8.60E-04 | 2.31E-02 |
| regulation of biological quality | GO:0065008 | 2820 | 24 | 7.39 | + | 3.25 | 4.87E-08 | 6.98E-06 |
| response to chemical | GO:0042221 | 3447 | 29 | 9.04 | + | 3.21 | 8.86E-10 | 2.47E-07 |
| positive regulation of biological process | GO:0048518 | 4413 | 37 | 11.57 | + | 3.2 | 3.04E-13 | 4.80E-10 |
| negative regulation of metabolic process | GO:0009892 | 2074 | 17 | 5.44 | + | 3.13 | 1.49E-05 | 8.46E-04 |
| regulation of catalytic activity | GO:0050790 | 1717 | 14 | 4.5 | + | 3.11 | 1.10E-04 | 4.42E-03 |
| regulation of cellular macromolecule biosynthetic process | GO:2000112 | 2849 | 23 | 7.47 | + | 3.08 | 2.83E-07 | 3.19E-05 |
| regulation of transcription, DNA-templated | GO:0006355 | 2541 | 20 | 6.66 | + | 3 | 3.59E-06 | 2.68E-04 |
| negative regulation of macromolecule metabolic process | GO:0010605 | 1908 | 15 | 5 | + | 3 | 8.85E-05 | 3.72E-03 |

| | | | | | | | | |
|---|---|---|---|---|---|---|---|---|
| regulation of biosynthetic process | GO:0009889 | 3059 | 24 | 8.02 | + | 2.99 | 2.30E-07 | 2.65E-05 |
| regulation of macromolecule biosynthetic process | GO:0010556 | 2935 | 23 | 7.7 | + | 2.99 | 4.84E-07 | 4.98E-05 |
| negative regulation of biological process | GO:0048519 | 3715 | 29 | 9.74 | + | 2.98 | 5.22E-09 | 1.06E-06 |
| negative regulation of cellular process | GO:0048523 | 3332 | 26 | 8.74 | + | 2.98 | 5.73E-08 | 7.68E-06 |
| regulation of RNA biosynthetic process | GO:2001141 | 2585 | 20 | 6.78 | + | 2.95 | 4.67E-06 | 3.34E-04 |
| regulation of nucleic acid-templated transcription | GO:1903506 | 2585 | 20 | 6.78 | + | 2.95 | 4.67E-06 | 3.32E-04 |
| signal transduction | GO:0007165 | 4150 | 32 | 10.88 | + | 2.94 | 5.41E-10 | 1.71E-07 |
| regulation of gene expression | GO:0010468 | 3246 | 25 | 8.51 | + | 2.94 | 1.58E-07 | 1.88E-05 |
| regulation of cellular biosynthetic process | GO:0031326 | 3023 | 23 | 7.93 | + | 2.9 | 8.21E-07 | 7.78E-05 |
| signaling | GO:0023052 | 4366 | 33 | 11.45 | + | 2.88 | 3.78E-10 | 1.38E-07 |
| regulation of RNA metabolic process | GO:0051252 | 2793 | 21 | 7.32 | + | 2.87 | 3.81E-06 | 2.83E-04 |

| | | | | | | | | |
|---|---|---|---|---|---|---|---|---|
| cell communication | GO:0007154 | 4463 | 33 | 11.7 | + | 2.82 | 6.86E-10 | 2.07E-07 |
| regulation of cellular metabolic process | GO:0031323 | 4654 | 34 | 12.2 | + | 2.79 | 3.96E-10 | 1.40E-07 |
| regulation of primary metabolic process | GO:0080090 | 4479 | 32 | 11.75 | + | 2.72 | 3.95E-09 | 8.51E-07 |
| regulation of nucleobase-containing compound metabolic process | GO:0019219 | 2965 | 21 | 7.78 | + | 2.7 | 9.81E-06 | 6.01E-04 |
| regulation of nitrogen compound metabolic process | GO:0051171 | 4387 | 31 | 11.5 | + | 2.69 | 1.16E-08 | 2.03E-06 |
| regulation of metabolic process | GO:0019222 | 4975 | 35 | 13.05 | + | 2.68 | 4.76E-10 | 1.57E-07 |
| regulation of cellular component organization | GO:0051128 | 1722 | 12 | 4.52 | + | 2.66 | 1.49E-03 | 3.50E-02 |
| regulation of macromolecule metabolic process | GO:0060255 | 4615 | 32 | 12.1 | + | 2.64 | 8.53E-09 | 1.62E-06 |
| cellular response to stimulus | GO:0051716 | 5401 | 35 | 14.16 | + | 2.47 | 4.92E-09 | 1.03E-06 |
| cellular protein modification process | GO:0006464 | 2208 | 14 | 5.79 | + | 2.42 | 1.40E-03 | 3.34E-02 |
| protein modification process | GO:0036211 | 2208 | 14 | 5.79 | + | 2.42 | 1.40E-03 | 3.33E-02 |

| | | | | | | | | |
|---|---|---|---|---|---|---|---|---|
| multicellular organismal process | GO:0032501 | 5130 | 32 | 13.45 | + | 2.38 | 1.22E-07 | 1.51E-05 |
| response to stimulus | GO:0050896 | 6436 | 37 | 16.88 | + | 2.19 | 3.48E-08 | 5.20E-06 |
| cellular macromolecule metabolic process | GO:0044260 | 3694 | 19 | 9.69 | + | 1.96 | 2.32E-03 | 4.91E-02 |
| regulation of biological process | GO:0050789 | 9715 | 48 | 25.48 | + | 1.88 | 8.09E-10 | 2.35E-07 |
| regulation of cellular process | GO:0050794 | 9152 | 45 | 24 | + | 1.88 | 1.38E-08 | 2.40E-06 |
| biological regulation | GO:0065007 | 10339 | 49 | 27.11 | + | 1.81 | 1.21E-09 | 3.18E-07 |
| cellular process | GO:0009987 | 12102 | 48 | 31.74 | + | 1.51 | 4.68E-06 | 3.31E-04 |
| biological_process | GO:0008150 | 15688 | 54 | 41.14 | + | 1.31 | 1.15E-05 | 6.84E-04 |
| Unclassified (UNCLASSIFIED) | | 5667 | 2 | 14.86 | - | 0.13 | 1.15E-05 | 6.86E-04 |
