## Supplementary material for "Equine synovial fluid small non-coding RNA signatures in early osteoarthritis": Primer sequences/assays used

Additional File 5

| Target | Acession number | Homologous to human sequence? | Primer sequence/Qiagen product |
|---|---|---|---|
| eca-mir-143 | MIMAT0013063 | Yes. hsa-miR-143-3p (MIMAT0000435) | Hs_miR-143_1 miScript Primer Assay |
| eca-mir-223 | MIMAT0013205 | Yes. hsa-miR-223-3p (MIMAT0000280) | Hs_miR-223_1 miScript Primer Assay |
| eca-mir-99a | MIMAT0013184 | Yes. Identical to hsa-miR-99a-5p (MIMAT0000097) | Hs_miR-99a_2 miScript Primer Assay |
| eca-mir-23b | MIMAT0013113 | No. | AUCACAUUGCCAGGGAUUACC |
| eca-let-7a-2 | MIMAT0012979 | Yes. hsa-let-7a-5p (MIMAT0000062) | Hs_let-7a_2 miScript Primer Assay |
| eca-mir-100 | MIMAT0012980 | Yes. hsa-miR-100-5p (MIMAT0000098) | Hs_miR-100_2 miScript Primer Assay |
| eca-mir-191a | MIMAT0013079 | Yes. hsa-miR-191-5p (MIMAT0000440) | Rn_miR-191_1 miScript Primer Assay |
| eca-mir-181a | MIMAT0013178 | Yes. hsa-miR-181a-5p (MIMAT0000256) | Hs_miR-181a_2 miScript Primer Assay |
| U6 | . | Yes. U6 | Hs-RNU6_2 miScript Primer Assay |
| snord96A | . | Yes. SNORD96a | MSC0076150 |
| snord13 | . | Yes. SNORD13 or U13 | MSC0076181 |
